# From Single-Cell Emergent Behaviors to Clinical Outcome: PTEN-driven Migratory Efficiency as a Potential New Vulnerability in Glioblastoma

**DOI:** 10.64898/2026.03.18.712310

**Authors:** Mariangela Morelli, Gianmarco Ferri, Francesca Lessi, Francesca Di Lorenzo, Sara Franceschi, Francesca Marchetto, Gabriele Tancreda, Tiziano Vadi, Francesco Sarnari, Tim Hohmann, Francesco Pieri, Carlo Gambacciani, Francesco Pasqualetti, Yawer Shaw, Jace Singh, Bruce West, Michele Menicagli, Manuel Giacomarra, Lucio Tonello, Paolo Aretini, Filippo Geraci, Aldo Pastore, Orazio Santo Santonocito, Anna Luisa Di Stefano, Paolo Grigolini, Luigi Palatella, Chiara Maria Mazzanti

## Abstract

**Background:** Glioblastoma (GB) is a highly aggressive brain tumor with a median survival of approximately 14 months, primarily due to its ability to infiltrate healthy brain tissue both as single cells and in collectives. A deeper understanding of GB cell motility, both individual and collective, is crucial for developing patient-specific therapies. We aimed to characterize migration in patient-derived GB cells using advanced modeling to identify stratification markers and therapeutic vulnerabilities.

**Methods:** We developed Single-Cell Behavior Live Imaging (ScBLI), an approach integrating live imaging with computational analysis, applied to 30 GB primary cell cultures. Trajectories and morphological features were tracked and analyzed. Diffusion Entropy Analysis (DEA) was applied to classify trajectories based on the Delta Scaling parameter (δ scaling). We evaluated functional responses correlating all findings with clinical outcomes and transcriptomic profiles.

**Results:** We analyzed 4,279 cell trajectories. Based on δ scaling (range 0.28–0.837), we defined three distinct motility groups: Low (L, δ scaling ≤0.5), Medium (M, 0.5 < δ scaling ≤ 0.7), and High (H, δ scaling >0.7). Functional assays demonstrated that Group H cells are more performant in both positive and negative chemotaxis. Clinically, the three groups showed a clear linear progression with patient survival: High δ scaling correlated with the shortest survival (poorer prognosis), while Low δ correlated with the longest survival, suggesting that structured motility drives invasiveness. Integrative multi-omic analysis, encompassing both exome and transcriptome profiling, demonstrated that these groups are defined by distinct molecular landscapes rather than poor behavioral traits. Moreover, exome data revealed that Group H is significantly enriched in PTEN alterations (75% vs. 8% in Group L), with PTEN gain-of-function (GoF) mutations exclusively restricted to this group (100% vs 0% in Group L). Notably, within our extended cohort (n=51) currently characterized by whole-exome sequencing, we observed that specific PTEN GoF mutations were associated with a significantly shorter survival compared to PTEN wild-type cases (median OS 6.4 vs 16.6 months; p=0.02), which typically harbor the canonical loss of chromosome 10q. A similar clinical trend was observed when comparing directly GoF carriers to patients with truncating (Ter) alterations (median OS 6.4 vs 14.3 months; p=0.09). Conversely, no survival difference was found between truncating (Ter) mutations and wild-type cases.

**Conclusion:** Our findings demonstrate for the first time that migratory efficiency, quantified through DEA, represents a powerful predictor of glioblastoma aggressiveness. Tumor cells adopting highly efficient exploration strategies are strongly associated with poor clinical outcomes and are characterized by distinct molecular signatures, notably PTEN gain-of-function alterations.

**Statement of significance:** Our multi-scale computational framework elucidates emergent behavioral phenotypes as pivotal drivers of glioblastoma progression. By demonstrating a correlation between enhanced migratory efficiency, PTEN gain-of-function, and significantly reduced overall survival, we establish a foundational paradigm for deciphering the emergent complexity governing tumor invasiveness.

## Introduction

Glioblastoma (GB) is the most common and aggressive primary brain tumor in adults, characterized by its devastating malignancy and notoriously poor prognosis, with a median survival of only 15-18 months despite aggressive surgical and chemoradiation therapies^1^. The defining clinical challenge that renders GB incurable is the diffuse infiltration of tumor cells into the surrounding healthy brain parenchyma, often far away from the primary tumor bulk^2,3^. This pervasive invasion makes complete surgical resection impossible and is the primary source of resistance to localized therapies, inevitably leading to recurrence^4^. The invasive phenotype of GB cells is driven by significant cellular and molecular heterogeneity^5^, with individual cells employing various migratory modes, ranging from single-cell (amoeboid or mesenchymal) to coordinated collective migration, along pre-existing structures like white matter tracts and perivascular spaces^6^. Crucially, the cellular proliferation-migration phenotype dichotomy also known as the “go-or-grow” dichotomy, is a central element in the behavior of several tumor types, where cells can choose between proliferation at the primary site (“grow phenotype”) or migration/metastasis towards distant sites (“go phenotype”)^7^. This strategic choice is highly plastic and deeply influenced by the tumor microenvironment (TME)^7^.

Traditional in vitro assessments of motility often rely on magnitude metrics, such as mean speed and total distance, or single parameters (e.g., displacement), which quantify the kinetic activity but are insufficient for capturing the underlying strategy or efficiency of migration^8^. GB migration involves a multi-faceted interplay of individual exploratory behaviors (e.g., persistence, directional change) and collective, coordinated movements (e.g., order, neighbor stability). A single metric fails to capture the intricate synergy between these two distinct but complementary levels of movement defined by a delicate balance and interconversion between individual cell exploration and collective group organization^9^.

To overcome these limitations and resolve the full spectrum of migratory dynamics, we innovatively applied Diffusion Entropy Analysis (DEA) to cell trajectories to quantitatively determine the characteristic δ scaling parameter (δ). DEA is a robust, scale-invariant method rooted in statistical physics that quantifies the type of motion^10^. Specifically, higher δ scaling (from >0.5 onwards) on denotes motion ranging from diffusive to super-diffusive (active, persistent, or directed), which is characteristic of highly efficient search strategies, often termed as Lévy walk movements ^11,12^ ^13^. Indeed, the finding that CD8+ T cell motility is well described by a generalized Lévy walk, a strategic search pattern similar to those employed by animal predators and other organisms, which allows T cells to find rare targets with greater efficiency than Brownian motion, underscores the critical relevance of analyzing such search patterns in tumoral cellular biology^14,15^. Understanding whether tumor cells also utilize such an efficient exploratory movement is crucial, as this inherent efficiency could represent a critical vulnerability to be targeted therapeutically.

Based on the δ scaling values, we intended to stratify samples in distinct motility groups. We hypothesized that the efficiency of cellular migration, quantified by δ scaling, would reveal a linear relationship with specific patient survival^11^. Additionally, we aimed to investigate the functional implications of these motility classes, and to identify distinct molecular signatures across the resulting behavioral clusters that could pinpoint new vulnerability targets.

## Results

### Glioblastoma Cell Motility Analysis via Single-Cell Trajectory Acquisition and Diffusion Entropy Analysis

Primary GB cultures were established from surgical specimens (**Table S1, Figure S1**) and analyzed together with the commercial normal human astrocyte line (NHA). Automated segmentation and tracking of individual cells produced a total of 4,213 single-cell trajectories across the cohort (**Figure 1, Supplementary Video S1, S2**), with 67–245 trajectories obtained per sample (**TableS2**). These trajectories were subsequently analyzed using DEA, a scaling-based approach that quantifies the statistical properties of cell motion by evaluating the Shannon entropy as a function of time (S(t)) (see Materials and Methods). The scaling exponent δ was estimated by fitting the entropy curve S(t) in the range of 20–100 timesteps (each timestep corresponding to one frame of the video). Robustness of the estimation was verified by repeating the analysis using slightly modified fitting intervals (**Figure 2a,b**).

**Figure 1:**
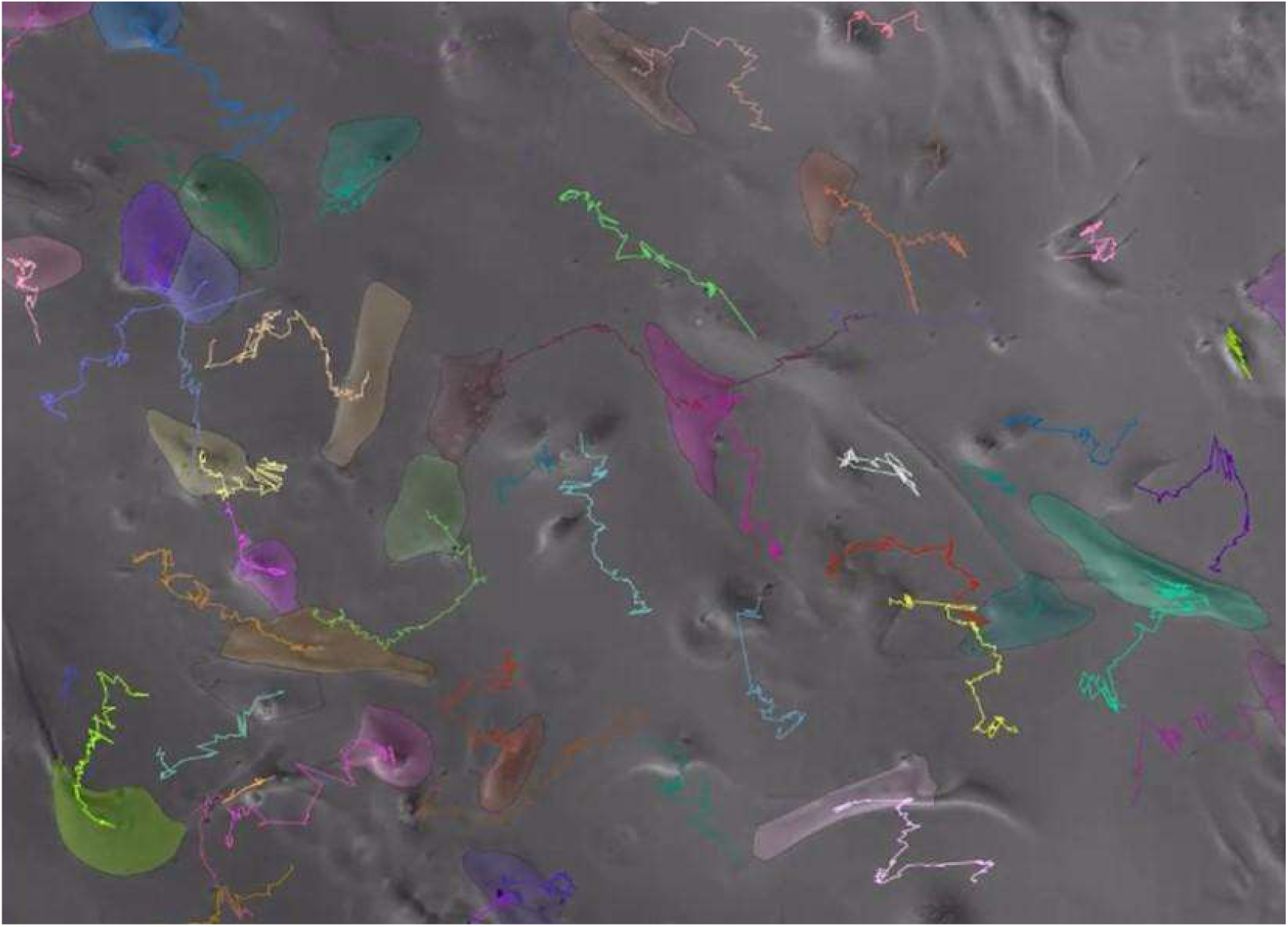
Automated Segmentation and Tracking of Patient-Derived Glioblastoma (GB) Cells. Representative single frame from a 72-hour live-cell microscopy video showing patient-derived GB cells. The cells were automatically segmented (Cellpose software). Each segmented cell (distinguished by color outlines in the original image, which would be visible in the figure is followed over time across the entire video duration. This automated tracking process generates continuous serial (x, y) coordinates for the centroid of each individual cell, which are then used to reconstruct the single-cell trajectories for subsequent dynamic analysis (Diffusion Entropy Analysis).

**Figure 2.**
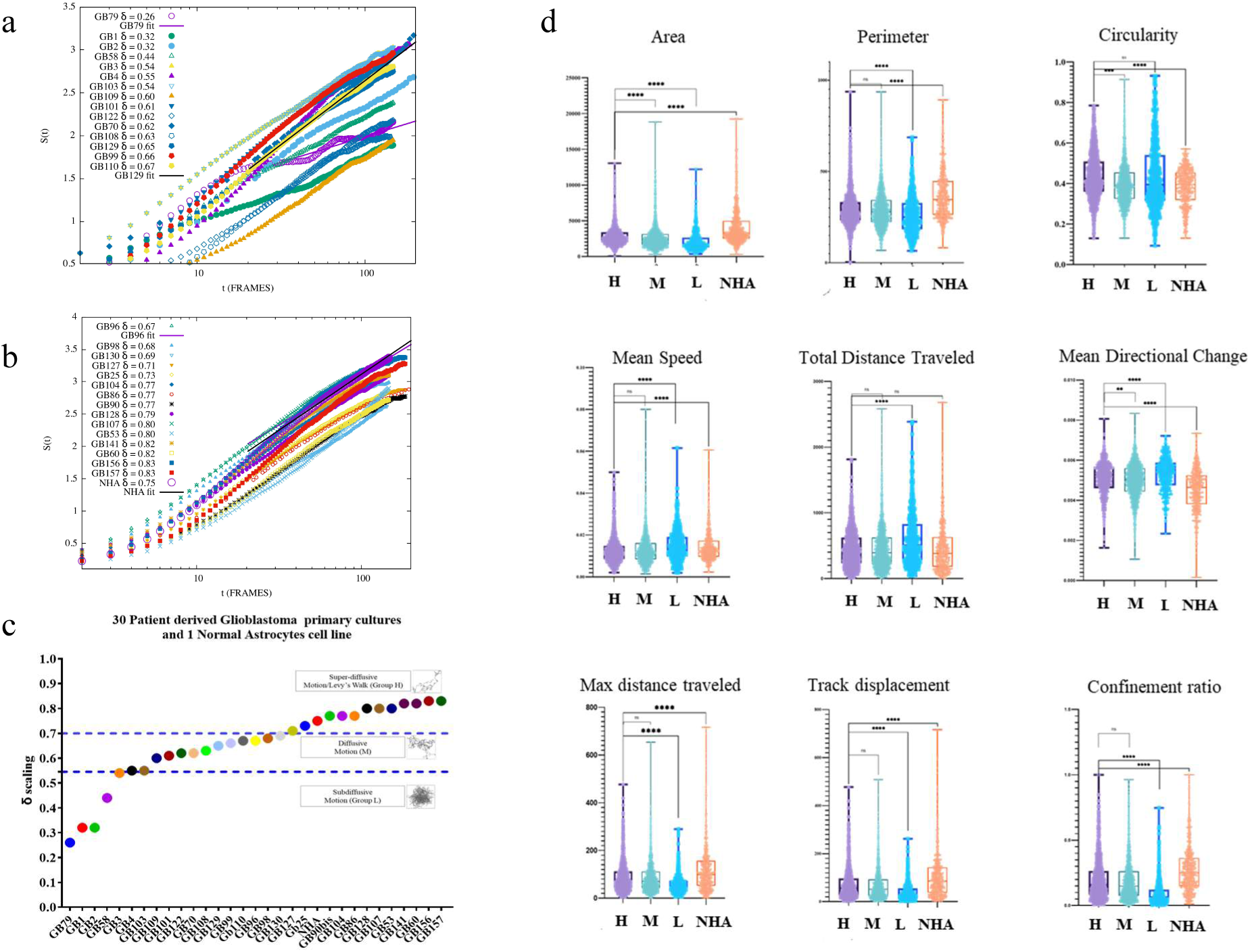
Diffusion Entropy Analysis (DEA) and scaling of cell motility. **(a)** Shannon entropy S(t) as a function of time (t) for GB patient-derived cell cultures and normal human astrocytes (NHA) has been applied. The straight lines in the log–linear plots (**a,b**) correspond to the best-fit for the lowest (GB79, GB96) and highest (GB129, NHA) scaling values within each specific group. In both panels, the slope of the curves in the log–linear plot represents the scaling exponent δ, obtained through a non-linear fit of S(t) over the interval of 20–100 timesteps. The linear behavior confirms that the diffusion process follows the scaling relation S(t)=A+δlog(t). **(c)** Distribution of the δ scaling parameter values, derived from the DEA, for 30 patient-derived GB primary cultures and one Normal Astrocytes Cell Line (NHA), with the cell lines sorted by increasing δ value. **(d)** Comparison of motility and morphological parameters are shown across δ scaling defined groups and NHA. Violin plots show the distribution of nine motility and morphological parameters, derived from the CellPose tracking analysis, across the three GB stratification groups based on the DEA δ scaling parameter (H, M, L), alongside NHA. Analysis of cell shape metrics reveals that the High δ scaling group demonstrates significantly reduced cell size (Area and Perimeter) and lower shape complexity (Circularity) compared to the Low δ scaling group and NHA. Furthermore, the data highlights a crucial, counter-intuitive relationship between migratory efficiency and kinematic magnitude: despite representing the most efficient, movement, the High δ scaling group exhibits significantly lower Mean Speed and Total Distance Traveled, compared to the Low δ scaling group. This indicates that strategic movement (Lévy-like) is optimized for efficient exploration (high Track Displacement) rather than simply maximizing speed or Total Distance Traveled. As expected the H group has significantly lower Mean Directional Change Rate and higher Confinement Ratio compared to the L group. Normal Astrocytes (HBA) exhibit distinct morphological characteristics compared to the GB groups. They possess significantly bigger Area and Perimeter but lower Circularity than all GB groups. Despite sharing the High δ Scaling efficiency with the most diffusive GB cells, HBA cells demonstrate significantly lower Mean Directional Change Rate and crucially, maintain significantly higher Mean Speed and Total Distance Traveled than all tumor groups.

The δ scaling parameter allows for the stratification of cellular movement into three distinct motility classes based on the nature of the diffusion observed (**Figure 2c**). These classes align with the theoretical frameworks of anomalous diffusion and Lévy statistics. **Figure 2c** displays the δ scaling values for 30 patient-derived GB primary cultures and NHA, sorted by increasing δ scaling value. The dashed blue line at δ=0.55 separates the motion types in: 1) Subdiffusive Motion (δ≤0.55): this group (L) exhibits motion where cells are highly restricted, characterized by dense, tangled trajectories indicative of a random walk with high probability of short steps and frequent directional changes. This motion is less efficient for long-range exploration; 2) Diffusive Motion (δ>0.5): this group (M) represents classical Brownian motion, where the mean square displacement increases linearly with time. This motion type is typically observed in cells showing standard, undirected random movement. Cell lines in this region span from approximately δ>0.55 to δ≤0.70; 3) Super-diffusive Motion (δ>0.70): this group (H) is characterized by efficient, organized movement, often termed as Lévy Walk. This strategic motion incorporates longer, persistent movement phases interspersed with random turns, a characteristic strategy of efficient search and invasive capability revealing a form of emergent “intelligence” at the single-cell level. This “intelligence” is hypothesized to arise from the inherent choices cells make in their movement, which are essential for GB invasion.

### Quantitative Analysis of Motility and Morphological Parameters

Quantitative descriptors of cell motility and morphology were extracted from the segmented trajectories (see Materials and Methods). The resulting dataset included a range of conventional and shape-related parameters, reported in supplementary data **Table S3**, including Area, Perimeter, Circularity, Track Displacement, Mean Speed, Total Distance Traveled, Max Distance Traveled, and Confinement Ratio. It is important to note that this analysis is based on individual cell parameters, which are then grouped by their respective motility class, H, M and L based on the δ scaling parameter. This stratification revealed significant differences in both motility and shape-related metrics across the three groups and in comparison with NHA (**Figure 2d**). Furthermore, the stratification of patient-derived GB cell coltures based on δ scaling reveals significant differences in these motility and shape metrics when comparing the H, M and L groups against each other and against NHA, as observed in **Figure 2d**. The number of individual cells analyzed, grouped by class of patient-derived cell line (H, M, and L δ scaling) and NHA, were exactly 1,505, 1,720, 512, and 565, respectively. Furthermore, this analysis highlights a crucial, counter-intuitive relationship between migratory efficiency (δ scaling) and kinematic magnitude. The High δ Scaling group exhibited higher efficiency and super-diffusive movement, suported by a significantly lower Mean Directional Change (MDC) and higher Confinement Ratio, as shown in **Figure 2d**. Paradoxically, despite this higher migratory efficiency, the H group also showed significantly lower Mean Speed and Total Distance Traveled compared to the L group. NHA cell lines exhibit distinct morphological characteristics compared to the GB groups. As expected for differentiated normal cells versus transformed tumor cells, the NHA cells possess a significantly bigger Area and Perimeter, but lower Circularity. This contrasts with the smaller, rounder morphology observed in the three GB cell groups. Regarding migratory dynamics, NHA cells share the motility signature of the H Scaling GB group, yet they show several key differences in magnitude. The NHA cells, characterized by high migratory efficiency, exhibit the lowest Mean Directional Change Rate among all groups, including the GB classes. Furthermore, unlike the tumor cells within the H group, the NHA cells demonstrate a significantly higher Mean Speed and Total Distance Traveled than all other groups. This indicates that while both the most efficient tumor cells and the normal astrocytes employ a strategic, Lévy-like exploration strategy (High δ Scaling), the normal cells maintain the highest kinetic magnitude (speed and distance).

### UMAP Clustering Reveals Behavioral Phenotypes and Inter-Tumoral Heterogeneity

To overcome the high dimensionality of the motility data set, the collected single-cell parameters, including Area, Perimeter, Circularity, Track Displacement, Mean Speed, Total Distance Traveled, Max Distance Traveled, Confinement Ratio, and Mean Directional Change Rate, were subjected to Uniform Manifold Approximation and Projection (UMAP) followed by K-means clustering^16^. This analysis incorporated the total of 4302 individual cells from the 30 patient-derived GB primary cultures and the NHA cell line. UMAP organized the data into three distinct behavioral groups, labeled as Cluster 0, Cluster 1, and Cluster 2 (**Figure 3a,b**). These clusters were defined by differential values across the input parameters, notably demonstrating large separation along Mean Directional Change Rate and Circularity. Investigating the composition of these clusters across individual cell lines, we observed significant inter-tumoral heterogeneity. **Figure 3c** illustrates that the percentage of cells belonging to V0 (Cluster 0), V1 (Cluster 1), and V2 (Cluster 2) varies substantially between the samples. Furthermore, the NHA line exhibited a cluster composition profile that was highly distinct from the majority of the other tumor cell lines. Crucially, comparison of cluster membership with the migratory efficiency measured by the δ scaling demonstrated a statistical significant correlation between Cluster V0 and δ scaling value (p=0.001). Conversely, the percentage of cells belonging to Cluster V2 was significantly and negatively correlated with the cell lines δ scaling value (p=0.04). This dual correlation establishes that the δ scaling of each cell line is directly mapped onto its underlying distribution of single-cell behavioral phenotypes (clusters) **(Figure 3d,e) (Table S4)**.

**Figure 3.**
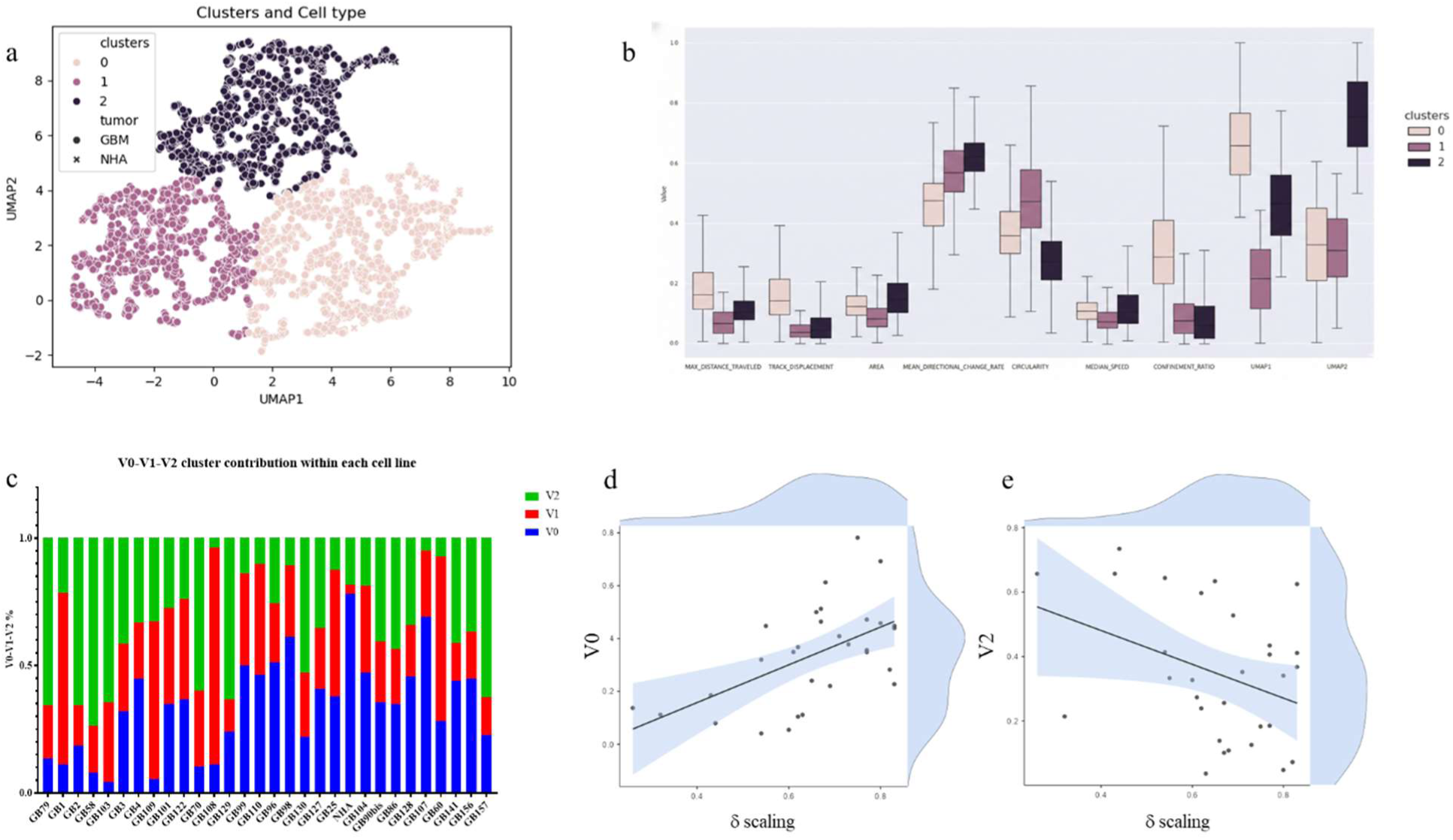
V0-V1-V2 cell subpopulation distributions within patient-derived GBM primary cells. **(a)** UMAP (Uniform Manifold Approximation and Projection) plot of n=4302 individual cells derived from 30 patient-specific GB primary cell lines and 1 healthy astrocyte (NHA) control line. K-means clustering (k=3, determined via the elbow method) identifies three distinct behavioral phenotypes: Cluster 0 (light pink), Cluster 1 (magenta), and Cluster 2 (dark purple). Individual cells are mapped based on a multidimensional feature set including circularity, area, and various kinematic tracking parameters. Solid circles indicate GB cells and solid crosses NHA cells; **(b)** Box-and-whisker plots illustrating the distribution of normalized feature values across the three identified clusters. Data represent the median, interquartile range (IQR), and whiskers extending to 1.5xIQR, with individual outliers indicated. Notable divergence between clusters is observed in parameters such as Mean Directional Change Rate, Circularity, and Confinement Ratio.**(c)** Stacked bar chart representing the relative proportions of V0 (blue), V1 (red), and V2 (green) cell states across 27 GB samples (GB1–GB27), with NHA (highlighted by the red box) serving as a baseline control**. (d,e)** Scatter plots with marginal density distributions demonstrating the correlation between the δ scaling metric and specific cell fractions, revealing a significant positive correlation with the V0 subpopulation (r=0.56, p=0.001) and a significant negative correlation with the V2 subpopulation (r=-0.36 p=0.04), suggesting that δ scaling is a key determinant of cellular state shifts and intratumoral heterogeneity.

### High δ Scaling Correlates with Chemo-Sensing Capability

To determine if the migratory efficiency described by the δ scaling parameter confers a functional advantage in environmental navigation, we conducted a modified Transwell chemotaxis assay (see Material and Methods). While traditional assays measure absolute invasive potential, our protocol was specifically designed to isolate the Sensing Index by decoupling gradient-directed movement from the intrinsic basal motility (M_bas_) of each line. To normalize for the inherent kinetic differences across the 14 patient-derived GB lines and the NHA control, we calculated the Sensing Index defined as (M_grad_-M_bas_)/M_bas_. This calculation mathematically isolates the specific response to external chemical stimuli by subtracting the spontaneous migratory component. The experimental design consisted of three distinct conditions: 1) In condition A (Basal Migration, M_bas_), as shown in **Figure 4a**, the nutrients (“food”) were provided in both the upper and lower chambers to eliminate chemical gradients, thereby measuring the stochastic ability of cells to cross the membrane without external stimuli; 2) in condition B (Positive Chemotaxis, M_grad_), as shown in **Figure 4b**, a nutrient gradient was established by placing food (FBS) solely in the lower chamber to measure the total migration driven by attractant-sensing; 3) in condition C (Negative Chemotaxis, M_grad_), as shown in **Figure 4c**, a gradient of the chemotherapeutic agent Temozolomide (TMZ) was established to quantify negative chemotaxis or a directional migration away from a toxic insult placed in the upper chamber..

**Figure 4.**
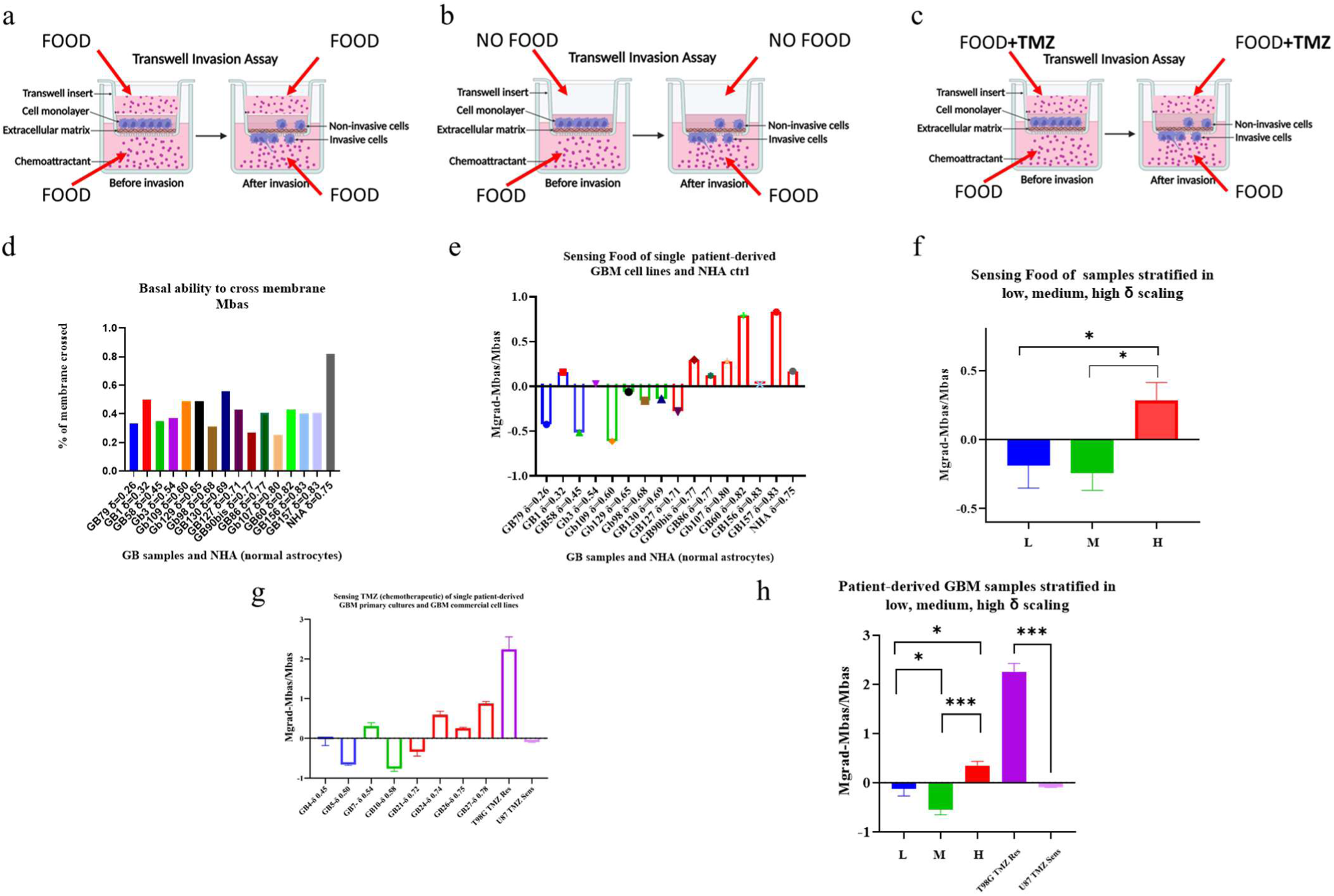
High δ scaling correlates with enhanced nutrient sensing and chemotherapeutic avoidance. **(a–c)** Schematic representation of the three experimental conditions for the modified Transwell Invasion Assay: **(a)** Condition A (M_bas_), no-gradient control; **(b)** Condition B (M_grad_), nutrient chemoattractant gradient; **(c)** Condition C (M_grad_), TMZ chemorepellent gradient. **(d)** Quantification of basal migratory capacity (Mbas) for 13 patient-derived GB lines and one NHA control. **(e,f)** Analysis of nutrient sensing. Left panel: Individual Sensing Index values; Right panel: Data stratified by δ scaling class, demonstrating significantly increased sensing in high δ cell cultures (*p<0.05). **(g,h)** Analysis of TMZ-avoidance (negative chemiotaxis). Left panel: Individual Sensing Index values including commercial controls. Right panel: Stratified analysis showing that high δ cell cultures exhibit significantly higher avoidance efficiency. Notably, TMZ-resistant control lines (T98G) show a significantly greater capacity to avoid the drug compared to TMZ-sensitive lines (U87 MG) (***p=0.001). Statistical significance determined via One-way ANOVA with post-hoc comparison tests.

As shown in **Figure 4d**, basal migratory capacity varied not significantly across the cohort. However, when normalized for sensing (**Figure 4e, g**), stratification by δ scaling revealed that H GB cell cultures possess a superior capacity to interpret environmental cues. High δ scaling cell cultures demonstrated significantly greater positive chemotaxis toward nutrients compared to low δ scaling groups (p<0.05) (**Figure 4f**). Crucially, in the negative chemotaxis assay, the inclusion of commercial controls revealed that TMZ-resistant high δ scaling lines (T98G) (data not shown) exhibit a significantly higher migratory capacity to avoid TMZ compared to TMZ-sensitive low δ scaling lines (data not shown) (U87) (p=0.001) (**Figure 4h**). This enhanced negative chemiotaxis in TMZ-resistant T98G models mirrors the behavior of high δ GBM patient-derived cell cultures, which also displayed significantly higher avoidance efficiency than lower scaling groups (p=0.01). These findings indicate that high δ scaling identifies a behavioral phenotype characterized by optimized navigational efficiency and active therapeutic evasion (**Table S5**).

### High δ Scaling Correlates with proliferation kinetics

We analyzed the proliferation kinetics of ten cell lines, including NHA and multiple GB-derived lines, across three seeding densities (500, 1500, and 3000 cells per well) over a 72-hour period. Cell proliferation was assessed using a Real-Time Glow luminescence assay.To quantitatively compare proliferative performance across cell lines and experimental conditions, we calculated the Area Under the Curve (AUC) of the normalized luminescence profiles (**Table S6**). The AUC^17^ was used as an integrated proxy of cumulative proliferative output, allowing assessment of growth dynamics across the entire observation window rather than relying on endpoint measurements. Correlation analysis between AUC values and the migration efficiency parameter (δ scaling) revealed a clear dependence on seeding density (**Figure 5a**). At the lowest density (500 cells per well), no association was observed between δ scaling and AUC values (r=0.12, p=0.742). A positive, albeit non-significant, trend emerged at intermediate density (1500 cells; r=0.53, p=0.112). In contrast, at the highest density (3000 cells per well), a significant positive correlation was detected (r=0.70, p=0.025), indicating that higher migration efficiency was associated with increased cumulative proliferative output under crowded conditions. Consistently, cell lines characterized by superdiffusive migration behavior (higher δ scaling values) maintained higher luminescence-derived growth trajectories at high density, whereas hypodiffusive lines exhibited earlier growth deceleration and reduced cumulative expansion (**Figure 5b**). These differences reflect sustained effects across the full-time course rather than transient variations at individual time points, highlighting a density-dependent coupling between migratory dynamics and proliferative fitness.

**Figure 5.**
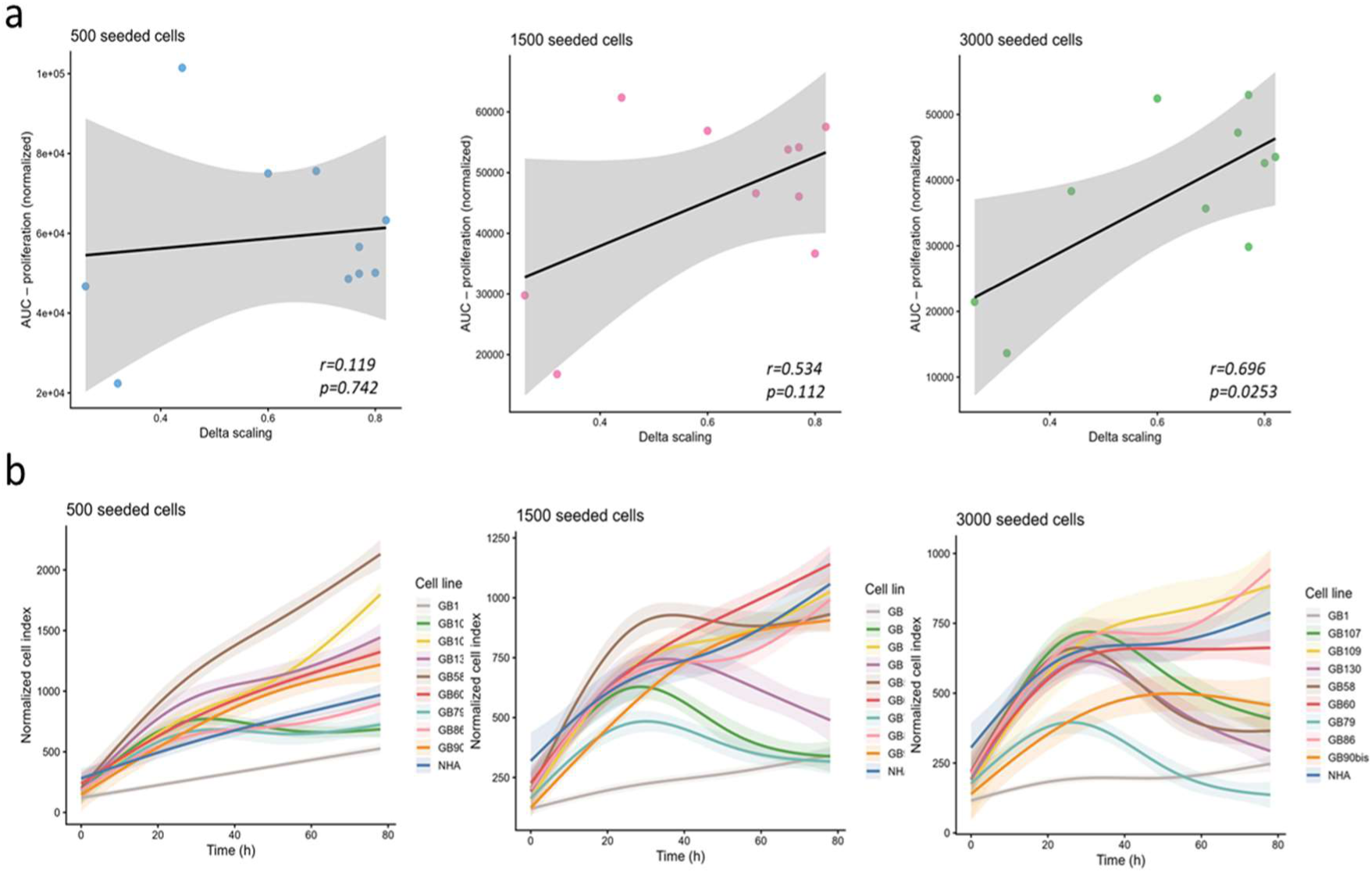
Proliferation kinetics and relationship with migration efficiency (δ scaling). **(a)** Correlation analysis between cumulative proliferative output and migration efficiency across three seeding densities. For each cell line, proliferation was quantified as the Area Under the Curve (AUC) of normalized Real-Time Glow luminescence profiles recorded over 72 hours. Scatter plots show AUC values plotted against the corresponding δ scaling parameter, an indicator of migration dynamics, for cells seeded at 500, 1500, or 3000 cells per well. Solid lines represent linear regression fits with 95% confidence intervals. **(b)** Proliferation kinetics of individual cell lines at increasing seeding densities. Curves represent smoothed growth trajectories derived from normalized luminescence measurements collected at multiple time points over 72 hours. For each cell line and density, proliferation profiles were modeled using generalized additive models (GAM), and shaded areas indicate the 95% confidence intervals of the fitted curves. At high seeding density, cell lines exhibiting superdiffusive migration behavior (higher δ scaling values) display sustained proliferative activity, whereas hypodiffusive lines show earlier growth deceleration.

### Integrating Clinical Outcomes and Redefine δ Scaling Stratification

To translate our in vitro findings into a clinically relevant framework, we conducted a retrospective survival analysis on a cohort of 24 patients diagnosed with GB IDH-wildtype, from whom the primary cultures were derived. Initially, we implemented a standard stratification strategy based on the three pre-established levels of migration efficiency: Low (L), Medium (M), and High (H) δ scaling. The Kaplan-Meier estimator revealed a striking and statistically significant correlation between these migration-based cohorts and Overall Survival (OS) (Log-rank p=0.009) as shown in **Figure 7a**. Patients characterized by a superdiffusive phenotype (High δ scaling, n=10) exhibited the shortest survival times, with a median OS of only 5.28 months. In contrast, the Medium group (n=10) and the Low group (n=4) demonstrated markedly improved outcomes, with average OS values of 17.4 and 27.0 months, respectively (**Figure 6a**). To further enhance the prognostic resolution of this parameter, we employed maximally selected log-rank statistic. This analysis yielded an optimal δ scaling cut-off of 0.670. Based on this novel threshold, the cohort was re-stratified into two statistically distinct groups: LowC (δ 0.670, n=11) and HighC (δ>0.670, n=13). This binary classification resulted in a more statistically significant difference (*p*=0.0003) (**Figure 6b**). Notably, the HighC group exhibited a markedly reduced median OS of 6 months, while the LowC group reached a median OS of 28.6 months. The sharp divergence of the Kaplan-Meier curves at the 0.670 cut-off underscores the superior predictive power of this optimized parameter. To move beyond categorical grouping and explore the continuous functional form of this relationship, the association between migration dynamics and OS was evaluated using Cox proportional hazards models incorporating spline terms for δ scaling. This analysis revealed a significant non-linear relationship between migration efficiency and patient outcome (likelihood ratio test, p=0.02; Wald test, p=0.03; concordance index=0.75). Spline-derived hazard ratios indicated that higher δ scaling values, corresponding to superdiffusive migration behavior, were associated with an increased risk of death, whereas lower δ values were associated with more favorable survival (**Figure 6c**).

**Figure 6.**
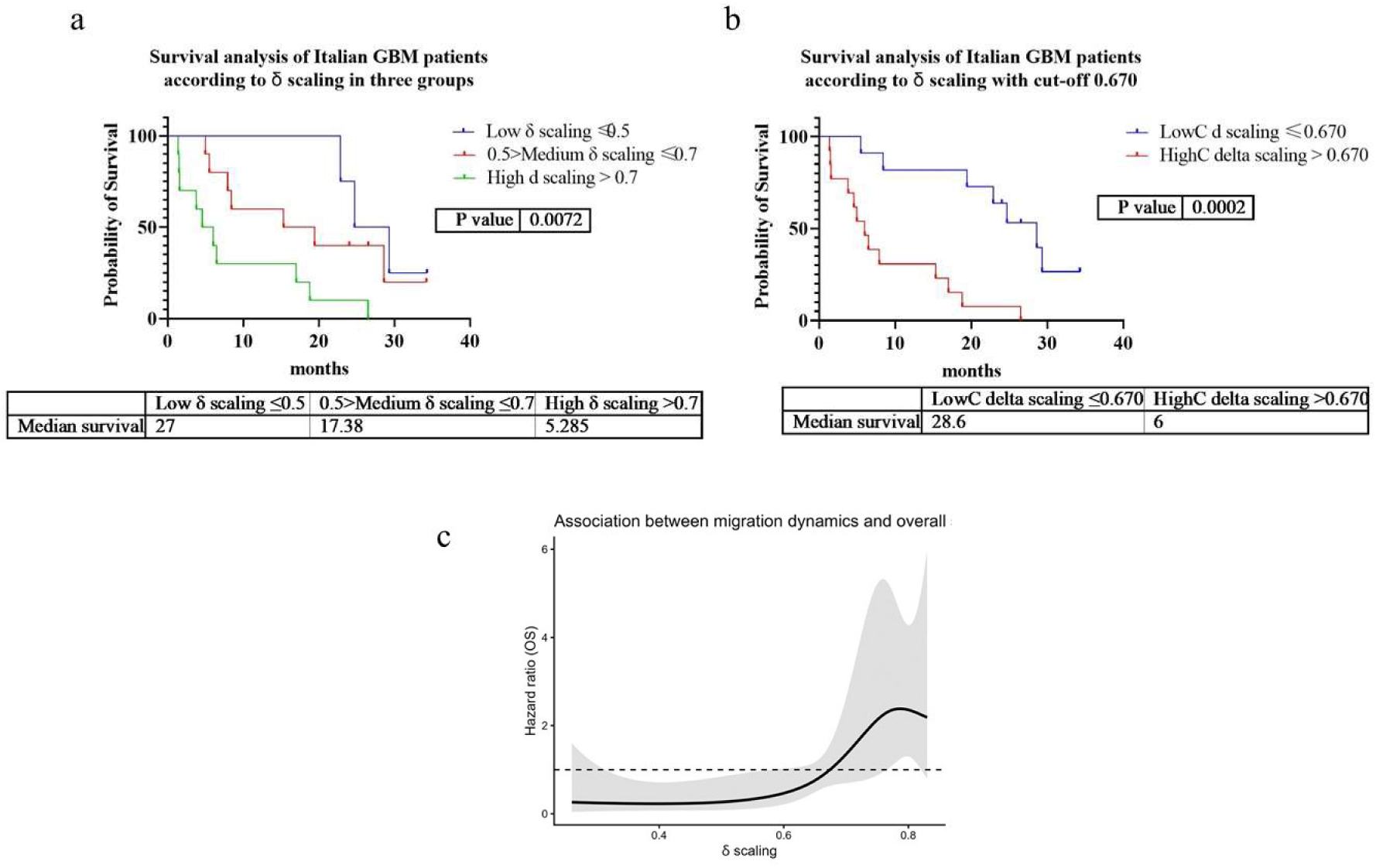
Survival stratification of GB IDH-wildtype patients based on migration efficiency. **(a)** Kaplan-Meier survival curves for the initial three-tier stratification such as Low, Medium, and High δ scaling (n=4, n=10, n=10 respectively). A significant correlation is observed between higher migration efficiency and reduced Overall Survival (OS) (Log-rank test, p=0.007). **(b)** Refined survival analysis using a maximally selected log-rank statistic cut-off of 0.670. The cohort (n=24) is divided into LowC (n=11) and HighC (n=13) groups. The optimized δ scaling threshold identifies two new patients populations with highly divergent clinical outcomes, where the HighC group exhibits significantly poorer prognosis (p=0.0002). **(c)** Spline-based Cox proportional hazards analysis illustrating the relationship between δ scaling and OS. The solid line represents the estimated hazard ratio as a function of δ scaling, while the shaded area indicates the 95% confidence interval. The dashed horizontal line denotes a hazard ratio of 1. Higher δ scaling values, indicative of superdiffusive migration dynamics, are associated with increased mortality risk. Confidence intervals widen at the extremes due to limited sample size.

**Figure 7.**
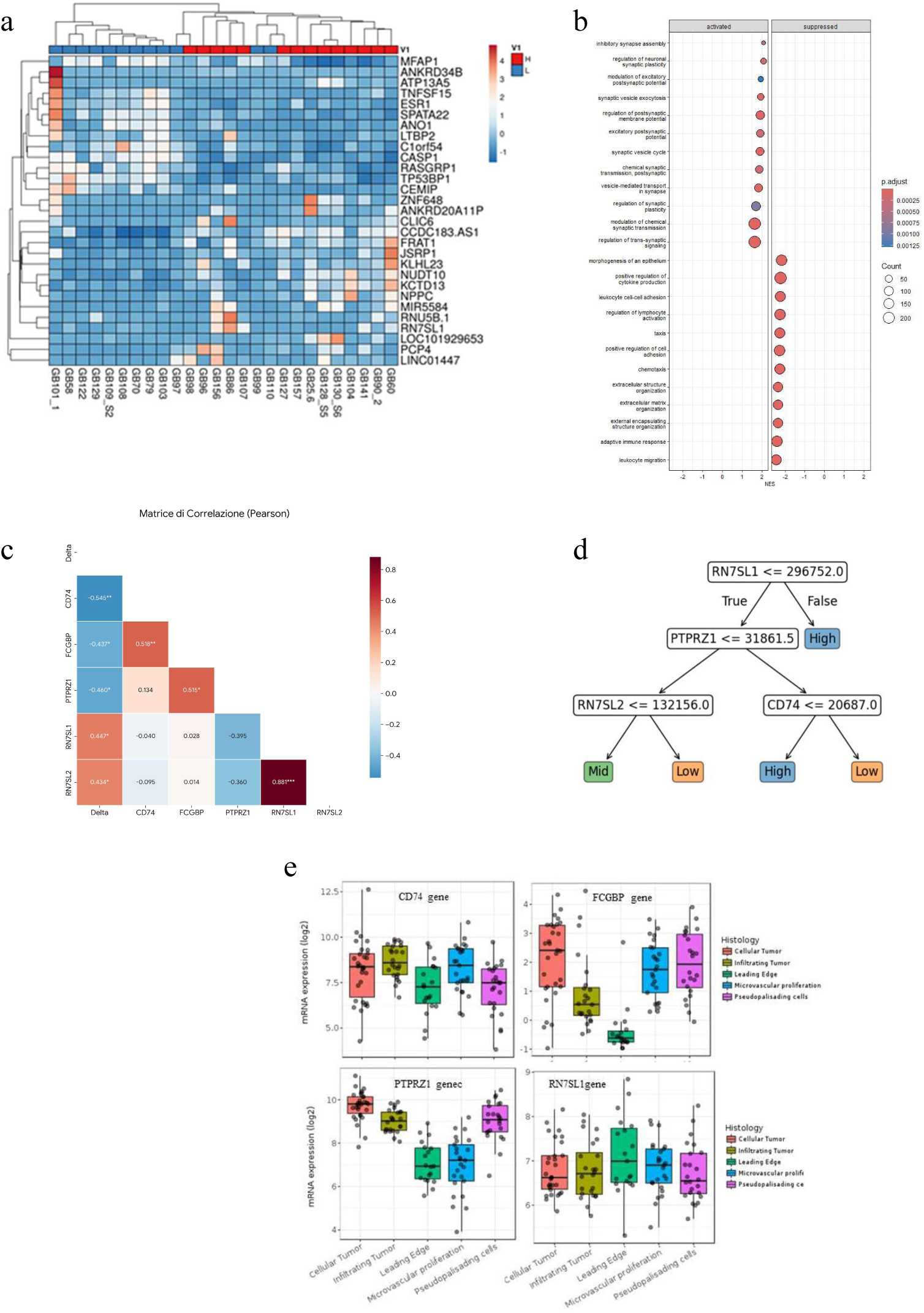
Multi-omic Characterization of the GB Migratory Phenotype. **(a)** Heatmap of Whole Transcriptome Analysis (WTA) displaying top 30 Differentially Expressed Genes (DEGs) between High (n=14) and Low (n=12) migratory cohorts, stratified by the δ scaling threshold of 0.670. **(b)** Gene Set Enrichment Analysis (GSEA) of identified DEGs, highlighting the significant involvement of these genes in cell adhesion and migration pathways. **(c)** The heatmap displays the Pearson correlation coefficients (r) for five selected genes (CD74, FCGBP, PTPRZ1, RN7SL1, and RN7SL2) linearly statistically correlated to the δ scaling parameter. The color scale indicates the strength and direction of the associations, ranging from dark blue (strong negative correlation) to dark red (strong positive correlation). Statistical significance levels are indicated by asterisks: *p<0.05, **p=0.01, ***p=0.001. **(d)** Contribution-based stratification model, enabling systematic sample categorization into Low, Medium, or High migratory phenotypes based on prioritized gene expression hierarchies and diffusion parameters. **(e)** Spatial validation utilizing the IVY GAP database: Gene expression signatures associated with high δ scaling are significantly enriched in the Leading Edge (invasive front) compared to central tumor zones.

### Comparative Transcriptomic and Mutational Profiling of HighC and LowC Migratory Phenotypes

To elucidate the molecular underpinnings of the observed migratory phenotypes, we performed a multi-omic characterization of patient samples, stratified into High and Low groups based on the previously identified δ scaling threshold of 0.670. Whole Transcriptome Analysis (WTA) revealed a molecular signature of Differentially Expressed Genes (DEGs) between the High (n=14) and Low (n=12) cohorts as shon by the Heatmap analysis (**Figure 7a**). Specifically, 8 genes were found to be significantly differentially expressed between the two groups with an adjusted pvalue<0.05, with 3 genes upregulated and 5 downregulated in the High group compared to the Low group (**Table S7**). Gene Set Enrichment Analysis (GSEA^18^) demonstrated that these DEGs are primarily involved in pathways related to cell migration, chemotaxis, adhesion and signaling (**Figure 7b**).

Beyond discrete group comparisons, we implemented a computational framework to treat δ scaling as a continuous quantitative trait, enabling the identification of genes whose expression levels correlate with migratory efficiency (see Materials and Methods). Using a contribution-based analytical approach, we identified a set of genes whose weighted contribution significantly influences the δ-scaling parameter (**Table S8**). Correlation analysis revealed structured relationships among these genes (**Figure 7c**), suggesting coordinated transcriptional programs associated with tumor cell motility. To further explore how these gene contributions collectively stratify the samples, we constructed a classification graph based on gene contribution weights (**Figure 7d**). This model illustrates how combinations of gene weights progressively separates samples into distinct groups characterized by different δ-scaling levels (e.g., low, intermediate, and high).

To investigate whether these transcriptional signals have spatial relevance within tumor tissues, we interrogated the Ivy Glioblastoma Atlas Project (IVY GAP) database. This analysis revealed that several genes associated with higher δ-scaling values show enriched expression in the Leading Edge region of glioblastoma samples (**Figure 7e**). The leading edge is the tumor compartment characterized by active invasion and parenchymal infiltration, representing the anatomical region where tumor cell migration is most prominent. Importantly, the spatial enrichment of these genes in the invasive front provides independent biological support for our computational stratification, linking the in vitro migratory phenotype quantified by δ scaling with in vivo spatial organization of tumor invasion.

### Exomic Profiling and PTEN Mutational Status

Finally, we performed Whole Exome Sequencing (WES) on the High and Low cohorts (12 vs 12). Matched blood samples were used to exclude germline variants. The analysis revealed several genes that were differentially mutated between the two groups, as illustrated in the oncoplot showing the top 20 mutated genes (**Figure 8a, Table S9**). Surprisingly, the analysis highlighted a highly significant divergence in the mutational status of the PTEN gene (**Figure 8b**). Specifically, PTEN mutations were identified in 9 out of 12 samples (75%) in the High δ scaling group, whereas only 1 out of 12 samples (8%) in the Low group harbored such alterations. This highly significant enrichment (p=0.03) suggests that an altered PTEN mutational state may be a key genomic driver of the superdiffusive and highly infiltrative phenotype in GB. **Figure 8b** details the specific types of mutations identified across the samples. As shown, the single mutation in the LowC group is a truncating mutation (Ter) occurring very early in the sequence, outside of the critical phosphatase or C2 membrane-binding domains. In contrast, missense mutations (GoF) are exclusively (100%) present in the HighC. Ter mutations are represented by three cases: one terminal mutation (suggesting a potentially functional protein) and two earlier mutations located specifically within the phosphatase domain (**Table S1**).

**Figure 8.**
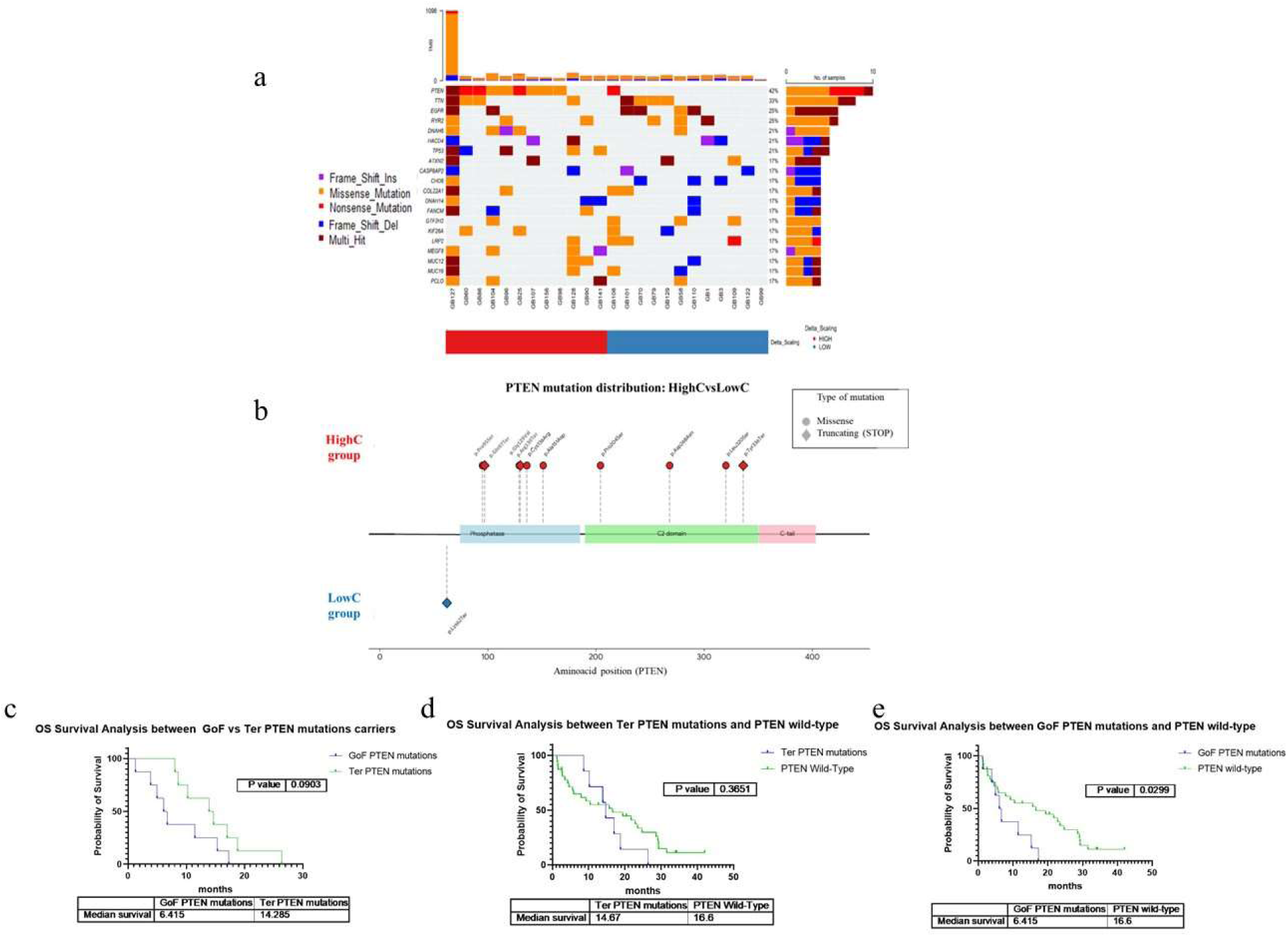
Clinical impact of PTEN mutation types on patient survival. **(a)** Mutational profiling via Whole Exome Sequencing (WES) of High and Low cohorts (n=12 vs n=12): The oncoplot illustrates the top 20 most frequently mutated genes. **(b)** Detailed mutational status of PTEN: a highly significant enrichment of mutations is observed in the High group (9/12) compared to the Low group (1/12). Specific mapping to functional domains (Phosphatase and C2) reveals a prevalence of missense mutations within the catalytic domain in the superdiffusive phenotype, whereas the Low group exhibits 1 early truncating mutation. **(c-e)** The comparison between GoF vs. Ter PTEN mutations (c) revealed a clear trend toward shorter survival for GoF carriers. When comparing Ter PTEN mutations against PTEN wild-type (d) no statistical significance was reached. Conversely, the analysis of GoF PTEN mutations vs. PTEN wild-type (e) demonstrated a statistically significantly shorter survival for the GoF group.

To further evaluate the clinical relevance of these findings, the survival analysis was performed on an expanded cohort of patients for a total of 59 GB samples **(Table S1-S10)**. This analysis revealed that the specific type of PTEN mutation significantly influences patient outcomes **(Figure 8c-e).** The comparison between GoF vs. Ter PTEN mutations (**Figure 8c**) revealed a clear trend toward shorter survival for GoF carriers (median 6.4 vs. 14.2 months; p=0.09). When comparing Ter PTEN mutations against PTEN wild-type (**Figure 8d**), no statistical significance was reached (median 14.67 vs. 16.6 months; p= 0.36). Conversely, the analysis of GoF PTEN mutations vs. PTEN wild-type (**Figure 8e**) demonstrated a statistically significantly shorter survival for the GoF group (median 6.415 vs. 16.6 months; p=0.02). Overall, these data validate the biological divergence of the H-group, confirming that PTEN gain-of-function alterations drive a significantly more aggressive clinical behavior compared to both loss-of-function (Ter) and wild-type configurations.

## Discussion

The relentless recurrence of GB is fundamentally rooted in its extraordinary capacity for parenchymal infiltration^19–21^. While surgical resection and radiotherapy can effectively debulk the primary tumor mass, the inability to control the “stealth” population of infiltrating cells, which migrate far beyond the contrast-enhancing margins, inevitably leads to fatal relapse. Despite decades of research, GB has never been stratified based on the biophysical strategies of these individual invaders. Our study fills this gap by demonstrating that the migratory efficiency, quantified through the δ scaling parameter, serves as a superior predictor of clinical outcome and a mirror of the underlying molecular landscape. The interpretation of our data suggests that the clinical aggressiveness of GB is not merely a function of kinetic speed, but rather a manifestation of a highly efficient navigation strategy quantified through the scaling parameter δ. Initially, our high-resolution single-cell tracking (**Figure 1**) revealed significant inter-tumoral heterogeneity (**Figure 2c**). By applying Diffusion Entropy Analysis (DEA)^10^, we categorized these diverse patterns into three distinct motility classes: Low (L, δ ≤0.55), Medium (M, 0.55 < δ ≤0.70), and High (H, δ > 0.70) (**Figure 2c**). This initial stratification demonstrated that while Group L cells exhibit localized and disordered movement, Group H cells adopt Lévy Walk dynamics^12^. This super-diffusive behavior, as shown in the scaling of Shannon entropy (**Figure 2a,b**), allows these “strategic navigators” to maximize environmental exploration while minimizing local redundancy. The normal human astrocyte (NHA) line aligns with this cohort, exhibiting a δ scaling value of approximately 0.75. This suggests that highly efficient motility is a component of physiological cellular dynamics and not exclusive to malignancy. Notably, however, certain GB cell lines also occupy this high-efficiency space, with Delta values reaching as high as 0.83. The Super-diffusive behavior observed in GBM cell lines is a key indicator of this adaptive migratory plasticity. Quantified by a δ scaling value greater than 0.55 (with values δ > 0.70 indicating particularly high migratory efficiency), this behavior suggests the frequent occurrence of Crucial Events (CE-s)^11^. In the 2D Lévy Walk model, these CE-s are interpreted not as simple random turns but as instantaneous corrections or abrupt, significant changes in the cell’s “flying direction”. This capability of the cells to execute and benefit from CE-s in their search strategy is a core aspect of their “intelligence”^11^. The complexity underlying this adaptive cellular migratory plasticity is theoretically linked to criticality, which suggests that the interaction among the internal components of the single cell drives this emergent behavior. A characteristic feature of criticality in these dynamics is that the waiting times between consecutive CE-s follow an Inverse Power Law (IPL) distribution with esponent μ. The DEA scaling index, δ, provides a direct connection to this underlying criticality. It is mathematically related to μ, through the theoretical scaling relation:

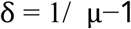

The manifestation of these precise scaling properties strongly suggests that single GBM cells possess a form of criticality-induced intelligence, affording them remarkable “adaptation to a changing environment” and the capacity for problem-solving necessary for highly invasive behavior ^10,11^ Interestingly, the quantitative analysis of motility and morphological parameters (**Figure 2d**) highlights a counter-intuitive relationship: despite their higher efficiency, High δ cells exhibit lower mean speed and total distance traveled compared to the Low group, suggesting that Lévy-like motion is optimized for exploration (high displacement) rather than simply maximizing speed or total movement distance^13^.

To further investigate the complexity of these dynamics and overcome the high dimensionality of the motility dataset, we applied UMAP and K-means clustering ^16^ to the single-cell parameters of over 4,300 cells (**Figure 3a,b**). This approach effectively organized the data into three distinct behavioral phenotypes, or clusters, primarily separated by differences in Mean Directional Change Rate and Circularity. While significant inter-tumoral heterogeneity was observed across the 30 patient-derived lines (**Figure 3c**), a key finding was the mapping of the δ scaling parameter onto these underlying distributions. Specifically, we observed a significant positive correlation between the percentage of cells in Cluster V0 and the δ scaling value (p=0.001), while Cluster V2 showed a significant negative correlation (p=0.04) (**Figure 3d,e**). These results would seem to suggest that within primary cell cultures, there exist three distinct sub-populations, two of which appear to be directly linked to the calculated migratory efficiency. Rather than a monolithic behavior, the overall δ value of a patient’s cell line might reflect a specific shift in the balance between these cell behavioral states.

The high migratory efficiency described by the δ scaling parameter appears to confer a distinct functional advantage in environmental navigation, as evidenced by our modified Transwell chemotaxis assays. By utilizing a protocol designed to isolate a sensing index, decoupling gradient-directed movement from intrinsic basal motility (M_bas_), we observed that basal migratory capacity alone did not vary significantly across the cohort (**Figure 4d**). However, once normalized for sensing, stratification by δ scaling revealed that High δ GB cell cultures possess a superior capacity to interpret and respond to environmental cues (**Figure 4e,f**). These High δ lines demonstrated significantly greater positive chemotaxis toward nutrient gradients compared to the Low δ groups. More strikingly, this superior navigation extends to active therapeutic evasion. In our negative chemotaxis assays (**Figure 4c**), High δ cultures displayed a significantly higher avoidance efficiency when exposed to a Temozolomide (TMZ) gradient. This behavior closely mirrors that of an established TMZ-resistant control lines (T98G), which exhibited also a significantly higher capacity to migrate away from the drug compared to TMZ-sensitive models (U87) (**Figure 4g,h**). These functional findings would seem to indicate that High δ scaling identifies a behavioral phenotype characterized not just by movement, but by optimized navigational intelligence and the ability to proactively escape toxic insults^22^.

Integrating the observations on colony formation and growth dynamics further contextualizes the biological relevance of our single-cell migration analysis. While, (**Figure S2**) immortalized commercial cell lines consistently form dense, macroscopic clusters^23^, a behavior mirrored even by normal human astrocytes (NHA), our patient-derived primary GB cultures demonstrated a markedly reduced capacity for colony formation across all δ scaling groups. Instead, these GBM primary cells remain largely dispersed, favoring an infiltrative and individualistic growth phenotype. This finding provides a critical biological rationale for our methodology: the absence of traditional colony-forming behavior confirms that primary GB cells are best characterized through single-cell behavioral tracking rather than bulk metrics, which may lose the characteristic dynamics inherent to the disease during the immortalization process. Consequently, our findings demonstrate that patient-derived primary models provide a significantly more reliable and physiologically relevant platform for studying the discrete migratory patterns and emergent complexity that define GB invasion. The link between these individual migratory strategies and collective tumor fitness is further elucidated by our analysis of proliferation kinetics (**Figure 5**). By calculating the Area Under the Curve (AUC) of normalized luminescence profiles as an integrated proxy for cumulative proliferative output, we observed a clear dependence on cell density. At the highest seeding density (3000 cells per well), a significant positive correlation emerged between δ scaling and cumulative growth (r=0.70, p= 0.025), indicating that higher migratory efficiency is associated with increased proliferative fitness under crowded conditions. These findings suggest a functional link between migration strategies and fitness that is modulated by the degree of cellular crowding ^24^. At low densities, where spatial constraints and nutrient competition are negligible, proliferative capacity appears independent of the mode of displacement. In contrast, at higher densities, the emergence of a positive correlation between δ scaling and AUC implies that more efficient migration strategies, specifically the long-range displacements characteristic of Levy-like walks, may confer a proliferative advantage. This could be attributed to the ability of highly motile cells to more effectively navigate crowded environments, potentially delaying the onset of contact inhibition or optimizing spatial distribution to access resources^25^. Conversely, the hypodiffusive behavior observed in samples with lower δ scaling may lead to local “entrapment,” resulting in reduced expansion capacity as the environment becomes saturated.

The clinical relevance of our biophysical stratification is most powerfully evidenced by the overall survival (OS) data, demonstrating that a purely physical measurement of trajectory efficiency can distinguish patient outcomes with high statistical significance. Initially, a three-tier stratification (Low, Medium, and High δ) revealed a striking correlation with prognosis: patients with a superdiffusive phenotype (High δ) exhibited a median OS of only 5.28 months, compared to 17.38 and 27.0 months for the Medium and Low groups, respectively. By further integrating survival data through employing a data-driven optimization, we identified a precise clinical δ threshold of 0.670, effectively bifurcating the cohort into two populations with highly divergent clinical paths: a HighC group with a median OS of 6 months and a LowC group reaching 28.6 months (**Figure 6b**). Analysis via Cox proportional hazards models and spline curves (**Figure 6c**) reveals a critical detail: the relationship between migration efficiency and mortality risk is non-linear. The spline curve highlights a threshold-like behavior, suggesting the existence of a critical threshold beyond which the risk increases precipitously. This threshold marks the transition toward Lévy Walk behavior, indicating that once a certain threshold of migration efficiency is reached, the tumor seems to accelerates disease progression. These findings could suggest that superdiffusion is not merely a laboratory observation but a potential driver of clinical aggressiveness, making the δ parameter a prognostic indicator.

The molecular landscape underpinning these behavioral shifts was further elucidated through a multi-omic characterization, which initially utilized a standard differential expression gene (DEG) analysis based on our clinically optimized δ threshold of 0.670. Hierarchical clustering revealed a distinct segregation between High and Low cohorts (**Figure 7a,b**), while Gene Set Enrichment Analysis (GSEA) confirmed that the most significant transcripts are primarily associated with cell migration, chemotaxis, and adhesion pathways (**Figure 7c**). However, to move beyond the limitations of discrete group comparisons, we leveraged the fact that δ scaling is an inherently continuous quantitative variable. We implemented an innovative computational approach to identify “ δ Scaling Gene Contributors”, genes whose expression levels correlate linearly or non-linearly with migratory efficiency across the entire spectrum (**Table S6**). Among these, we identified five primary drivers (CD74, FCGBP, PTPRZ1, RN7SL1, and RN7SL2) with high statistical significance (**Figure 7d**). These contributors were subsequently integrated into a Decision Tree model, used as a discovery roadmap to hierarchically rank the most influential biological drivers and systematically categorize samples into their respective migratory phenotypes. The resulting transcriptional signature validates a transition from a stationary/proliferative state to an invasive/secretory one. Specifically, the downregulation of PTPRZ1 and CD74 in High δ cells is indicative of a “detachment” from the local matrix, a necessary requirement for long-range dispersal. PTPRZ1 (Protein Tyrosine Phosphatase Receptor Type Z1) is a known mediator of cell-matrix adhesion^26,27^; its reduction suggests a strategic detachment that allows cells to move away from the tumor core toward white matter tracts. Similarly, the decrease in CD74 expression indicates a shift away from the pro-inflammatory niche of the tumor core toward an invasive phenotype^28,29^. Simultaneously, the upregulation of RN7SL1 (7SL RNA) could represents a critical transcriptional switch^30^. As the core component of the Signal Recognition Particle (SRP), the significant transcriptional over-expression of RN7SL1 in Group H signifies a specialized secretory hyper-activation. We hypothesize that RN7SL1-mediated secretion, potentially via exosomes, could prime the distant brain microenvironment by facilitating the fluidification of the extracellular matrix^31^. To assess the spatial relevance of these findings, we interrogated the IVY Glioblastoma Atlas Project (IVY GAP) database^32^. This analysis confirmed that the gene expression signatures associated with High δ scaling are significantly enriched in the Leading Edge (LE) of the tumor tissue (**Figure 7e**). This spatial validation reinforces the link between our *in vitro* metrics and the *in vivo* invasive front, suggesting these cells are programmed to focus on the colonization of healthy tissue. These findings may provide a biophysical mechanism for the recent spatial transcriptomic observations by Migliozzi et al.2025^33^, who demonstrated that cancer cell plasticity is restrained by spatial clustering, whereas cellular dispersion triggers a shift toward highly aggressive, plastic archetypes (GPM state). Our results propose the Lévy Walk as a fundamental migratory strategy could enable this transition. While Group L cells might remain in a more clustered, predictable and vulnerable state, Group H cells may exploit super-diffusive dynamics to actively escape homotypic restraint, reaching the dispersed state described by Iavarone et al., thereby gaining the plasticity required to survive therapeutic insults. Finally, to investigate the genomic determinants underlying the diverse migratory strategies observed, we conducted a Whole Exome Sequencing (WES) analysis on the HighC and LowC cohorts; among the top twenty most frequently mutated genes, a striking and statistically significant divergence emerged regarding the mutational status of PTEN^34^. The analysis reveals a near-absolute correlation between the superdiffusive phenotype and PTEN alterations, identified in 75% of samples in the HighC group (9/12) compared to only 8% in the LowC group (1/12). In this context, the behavior of the High δ group would seem to suggest a distinctive application of the Knudson ‘two-hit’ model^35^. While the loss of one PTEN allele is a hallmark and nearly universal event in GB, our findings highlight a critical secondary event in high-efficiency samples: the remaining wild-type allele is not merely deleted, but specifically mutated. This ‘double-hit’ scenario likely results in a gain of function rather than a simple loss of activity. We hypothesize that whatever is left of the PTEN signaling machinery is reprogrammed by these missense mutations to act as an unfettered navigational engine. The nature of these mutations would seem to dictate the migratory fate: the single mutation found in the Low group is an early truncating alteration, which could be equated to a complete loss of the protein (Loss of Function), resulting in a high-frequency stochastic motility but restricted and non-directional movement. In contrast, mutations in the High group are predominantly missense alterations falling specifically within the catalytic phosphatase domain (e.g., the R130 hotspot) or occurring very late in the coding sequence, as reported in the work of Choi et al. (2021)^36^. These GoF mutations might trigger a non-canonical gain of function, inducing the mutant protein to localize to the cell periphery and co-localize with F-actin and Cdc42 at the leading edge. This molecular shift could provide the biophysical basis for the “hopping” phenomenon described by Yasui et al. (2014)^37^. In physiological states, PTEN utilizes positively charged residues in its C2 domain (specifically the alpha helix) to suppress excessive hopping and stabilize membrane binding; the loss of this regulated brake in missense-mutant cells might transform the movement into the ballistic Lévy Walk trajectories we observed, enabling long-range, persistent displacements. Finally, the literature suggests that these specific phenotypes driven by missense mutations, while being resistant to PI3K/Akt inhibitors (such as BKM120), maintain a sensitivity to microtubule inhibitors like Colchicine, offering a potential therapeutic vulnerability for the High δ patient subset. In conclusion, the δ parameter might act as a functional integrator of the genomic state, identifying the critical point where GB evolves into an unstoppable and efficient infiltrative system^38^.

### Conclusions

The infiltrative nature of GB represents a formidable challenge that transcends traditional oncological frameworks, necessitating a shift toward understanding the tumor as a dynamic complex system. By adopting a multi-scale systems biology approach, inspired by the principles of emergent properties and complex systems (Parisi, 2021), we have demonstrated that GBM progression is driven by the behavioral dynamics of its elementary components. Unlike commercial cell lines that favor collective aggregation, the patient-derived primary cells analyzed in this study exhibit a distinct preference for individual growth and migration. This behavior validates our single-cell tracking methodology in non-confluent conditions as a physiologically superior model for capturing the authentic infiltrative strategy of the disease. Crucially, the phenotypic stratification of our patient cohort into three subgroups based on these migratory patterns revealed a significant linear correlation with overall survival, a finding that was further corroborated by distinct gene expression and mutational profiles. These results suggest that the “emergent” physical spreading of GBM cells is not a stochastic process, but a direct manifestation of the underlying molecular landscape, providing a robust and reliable proxy for clinical outcomes. Our study presents a novel framework for understanding and prognosticating GB aggressiveness by focusing on the efficiency of cellular movement, quantified by the δ scaling exponent from DEA. It is noteworthy that normal astrocytes (NHA) exhibit high migratory efficiency while maintaining both high speed and total distance. This suggests that aggressive High δ tumor cells hijack a physiological, super-diffusive program, modeled as a Lévy walk, to navigate the brain parenchyma. This efficient search pattern mirrors the behavior of CD8+ T cells, which utilize generalized Lévy walks to find rare targets with greater efficiency than Brownian motion. While normal cells maintain high kinetic magnitude, the most aggressive GB cells appear to trade speed for persistence. Low δ tumor cells remain kinetically frenetic, moving fast but in a confused, near-random manner. This lack of directional coherence may render them more vulnerable and less invasive compared to the slower, yet highly efficient navigators of the High δ group. The most critical finding is the direct and highly significant correlation between High δ Scaling and poor patient survival, establishing δ scaling as a powerful, quantitative prognostic biomarker. The discovery of a genomic “double-hit” in the High δ cohort, where the ubiquitous loss of one PTEN allele is followed by a specific missense mutation, proposes a fundamental shift in our understanding of this tumor suppressor. These mutations can transform the classic tumor suppressor PTEN into an oncogene, creating an unregulated navigational mechanism through the disruption of membrane-hopping kinetics. Ultimately, while these findings suggest that the Lévy walk is the biophysical engine of the infiltrative phenomenon, further experiments are needed to fully circumscribe this biological phenomenon and determine if targeting the PTEN-microtubule axis can effectively dismantle the effective migratory efficiency of the disease. A critical open question remains whether this scaling behavior reflects an intrinsic molecular background, potentially selected through evolutionary pressure, that predisposes cells to specific migratory strategies, or if it emerges primarily as a mechanically-induced response to external constraints. Distinguishing between an adaptive, molecularly-primed trait and a purely emergent physical effect is essential to understanding the robust nature of these dynamics. Recent work by Scita et al. (2025)^39^ demonstrates that tumor collectives exhibit a profound mechanical memory, where physical deformations induce transcriptional reprogramming via ATF3 upregulation, ultimately enhancing invasive potential. Consistent with these findings, the high δ scaling in our GB models suggests that a specific molecular landscape may lower the threshold for mechanically-triggered transitions, shifting cells from a stabilized state toward a highly persistent, super-diffusive regime

## Materials and Methods

### HBA and GBM Primary Cell Cultures

Commercial Human Brain Astrocytes (HBA, Alphabioregen, Boston, Massachusetts, USA; Cat# HBMP-202) are seeded in Astrocyte-Growth Medium (Cat# 821-500) and maintained in an incubator at 37 °C and 5% CO₂. Primary GB cell cultures (n=30) were established from patient-derived tumor explants via mechanical dissociation using enzymatic components provided in the Brain Tumor Dissociation Kit (Myltenyi Biotech, Bergisch Gladbach, Germany), following the manufacturer’s instructions. Primary cells were maintained at 37 °C in a humidified incubator with 5% CO₂, in the cultured medium consisted of DMEM/F12 medium (Gibco, Carlsbad, CA, USA), supplemented with 10% FBS (Gibco, Carlsbad, CA, USA), 100 U/mL penicillin, and 0.1 mg/mL streptomycin (Gibco, Carlsbad, CA, USA), 1% Amphotericin B (Gibco, Carlsbad, CA, USA), 1% G-5 supplement (Gibco, Carlsbad, CA, USA). Culture medium for HBA and primary GBM cell cultures was replaced the day after seeding. Upon reaching confluency, cells were used for the subsequent experiments.

### Live-Cell Imaging and Time-Lapse Microscopy

Time-lapse video were performed on a Zeiss Axio Observer Z1 inverted microscope equipped with a 10x air objective and configured for phase-contrast imaging. The microscope was fitted with an incubation chamber to maintain a controlled environment of 37°C and 5% CO₂ throughout the acquisitions.For single-cell motility tracking, approximately 100,000 primary cells were seeded onto 35 mm glass-bottom dishes and allowed to adhere overnight before imaging. Time-lapse videos were recorded for 18 hours, with a total of 220 frames captured at a rate of one frame every 5 minutes. To increase the dataset, at least four time-lapse videos were simultaneously recorded from different fields of view on the same sample using a tiling acquisition mode.For the analysis of confluent cultures, cells were seeded at a higher density and grown until a confluent monolayer was formed. The dynamics of the confluent layer were then monitored for an extended period of 72 hours under the same microscopy and incubation conditions.

### Video Analysis and Data Extraction

Cell Segmentation and Tracking for Motility Analysis. Time-lapse videos for trajectory analysis were processed using a custom pipeline integrating Cellpose for cellular segmentation directly in TrackMate, the ImageJ plugin^40^. Specifically, cell segmentation was performed using the pre-trained “cyto2” model in Cellpose. The resulting segmented cell outlines were then used for tracking analysis, carried out using the Linear Assignment Problem (LAP) tracker algorithm present in TrackMate. Each identified trajectory was manually reviewed to ensure accuracy. From this analysis, parameters describing the morphology and dynamic traits of the imaged cells were obtained for each track.

### Statistical Analysis and δ Scaling Evaluation

The statistical method employed to evaluate the scaling behavior of cell diffusion is hereby described. The experimental data, representing the positions of the cells, are treated as 2D projections onto a plane of a 3D motion occurring within a fluid medium. Fluctuations along the vertical direction, defined by gravity, are assumed to be limited and are therefore neglected in this analysis.

The issue of scaling evaluation can be illustrated by considering a simplified model of one-dimensional walkers. These walkers are assumed to start from the origin, x=0, at time t=0. As time increases, the initial distribution, represented by a Dirac delta function at x=0 evolves into an increasingly broad cloud. The width of this distribution is proportional to t^ δ, where δ denotes the scaling index. To ensure an accurate evaluation of this scaling, a high number of trajectories is required. In principle, the scaling δ could be determined with precision from a single trajectory, provided it is sufficiently extended in time, by utilizing the moving window method.

The moving window method is based on partitioning a single long trajectory into multiple segments (windows) of the same temporal duration. In this framework, the start of each window is interpreted as the temporal origin (t=0), and the initial position of the walker within that window is set as the spatial origin (x=0), regardless of its absolute coordinates. The displacement is then calculated as the difference between the position of the walker at the end of the window and its position at the start. This procedure allows for the generation of multiple trajectories that are considered statistically independent, despite being derived from the same underlying stochastic process.

The data under study were generated by observing the motility of various distinct cells. In principle, the scaling δ could have been evaluated without the moving window method by treating each cell as a single data point. However, the duration of each individual cell trajectory was insufficient for a statistically robust scaling evaluation. Consequently, to capture the necessary statistics, moving windows generated across different cells were utilized.

In this 2D framework, the coordinates of the k-th cell at physical time τ are denoted as X_k(τ) and Y_k(τ). By applying the moving temporal window method, several diffusive trajectories ΔX_k,i(t), ΔY_k,i(t) are generated for different temporal window sizes, denoted by the symbol t. It is important to distinguish t, the window duration, from the absolute physical time τ. The displacements are defined as:

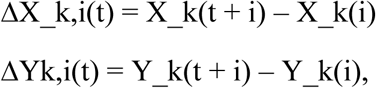

where 0 ≤ i < τ_max−t is the index indicating the starting position of the moving window as it slides throughout the entire sequence. By symmetry, the statistics of ΔX_k,i are assumed to be identical to those of ΔY_k,i. The total number of trajectories provided by this approach is determined as follows: assuming each cell sequence has a total duration of τ_max, for any temporal window of size t, there are 2Ncells(τmax − t) trajectory positions, denoted by the variable x.

The probability density function (PDF), p(x, t), is then computed. Although the number of available diffusive trajectories varies for different values of t, this is accounted for by normalizing p(x, t) such that the standard condition, Σ *p*(*x*, *t*) = 1 is satisfied for each t. If the diffusion follows a scaling prescription such that:

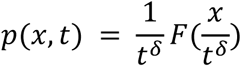

then the Shannon entropy is given by:

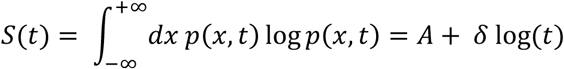

Consequently, the δ scaling parameter is obtained by calculating the slope of the function S(t) on a log-linear scale.

High-Dimensional Phenotypic Analysis: UMAP and Clustering on all Cellpose parameters Analyses were performed using Python (version 3.12). Parameters derived from Cellpose segmentation of both tumor and normal cells were used to generate a phenotypic characterization. Specifically: max_distance traveled, track displacement, area, mean_directional change rate, circularity, median_speed, and confinement_ratio were used as input features for the Uniform Manifold Approximation and Projection (UMAP) algorithm. A semi-supervised version of the UMAP algorithm was employed, which constrained the normal cells to cluster together in the projected two-dimensional space. This approach established a reference distribution for normal cells in the UMAP embedding, thereby enabling the identification of tumor cells with phenotypes similar to the normal control.

Following dimensionality reduction, the cells were grouped using spectral clustering (http://scikit-learn.org/stable/modules/generated/sklearn.cluster.SpectralClustering.html#sklearn.cluster.SpectralClustering), with the number of clusters set to three. For each patient, the percentage of cells belonging to each of the three clusters was calculated. This procedure resulted in each subject, including the healthy control, being described by a three-component vector representing these percentages.

Finally, these three new variables were used to cluster the patients via a k-means algorithm (https://scikit-learn.org/stable/modules/generated/sklearn.cluster.KMeans.html) with k set to 3. This final step allowed for the identification of phenotypically similar patients and, importantly, highlighted the subgroup whose cancer cells exhibited a phenotype similar to that of the healthy sample.

### Cell Migration Assay

The cell migration capacity was evaluated using polyethylene terephthalate (PET) inserts with a 5 μm pore size for 12-well plates (Sarstedt, Nümbrecht, Germany;). The cells were maintained in a humidified incubator until required, after which they were detached using 0,25% trypsin-EDTA (Gibco, Carlsbad, CA, USA) and counted using a cells counter (LUNA™ Automated Cell Counter, L10001, Logos Biosystems Inc). A total of 300,000 cells were seeded into the upper chamber of each insert. Three experimental conditions were tested: in the control condition, the cells were suspended in complete culture medium, and the same medium was added to the lower chamber; in the positive chemotaxis condition, the cells were suspended in a medium containing 1% FBS, while complete medium was added to the lower chamber; and in the TMZ treatment (Sigma-Aldrich, St. Louis, MO, USA) condition, the cells were suspended in a medium containing 250 µM TMZ, and complete medium was padded to the lower chamber. The plates were then incubated for 24 hours at 37 °C in a humidified atmosphere with 5% CO₂. After incubation, the migrated cells on the membranes were fixed with 4% paraformaldehyde (PFA, Life Technologies, Thermo Fisher Scientific, Italy) for 10 minutes at room temperature, followed by staining with crystal violet (0.1% crystal violet, 20% methanol, in water) for 15 minutes. The membranes were then thoroughly rinsed with PBS to remove excess stain. Each membrane was carefully detached from the plastic support by cutting around the edge with a sterile scalpel. The upper surface of the membrane, which contained the non-migrated cells, was gently wiped using a sterile cotton swab. Only the bottom side of the membrane, containing the migrated cells, was mounted on a microscope slide and scanned using the Aperio CS2 slide scanner (Leica Biosystems).

### Colony Assay

The colony formation assay was performed on HBA cells, three commercial and primary GBM cells. A total of 500, 1000 and 2000 cells per well were seeded in duplicate in 6-well plates and incubated for two weeks at 37 °C in a humidified atmosphere with 5% CO₂. Following incubation, the resulting colonies were fixed with 4% paraformaldehyde (PFA, Life Technologies, Thermo Fisher Scientific, Italy) for 10 minutes at room temperature, after which they were stained with crystal violet (0.1% crystal violet, 20% methanol, in water) for 15 minutes. The plates were then washed with distilled water to remove excess dye and air-dried prior to analysis.

### Proliferation assay and kinetic analysis

Cell proliferation was assessed using a Real-Time Glow luminescence assay (Promega Corporation, Madison, WI, USA). Cells were seeded in multiwell plates at three different densities (500, 1500, and 3000 cells per well) and monitored over a 72-hour period. Luminescence signals, proportional to metabolically active cell number, were recorded at multiple time points throughout the experiment. Raw luminescence values were normalized to enable comparison across cell lines and experimental conditions.

To quantitatively capture proliferation dynamics over time, growth curves were generated from normalized luminescence profiles. For each cell line and seeding density, replicate measurements were aggregated and analyzed to extract both kinetic and cumulative growth features. The Area Under the Curve (AUC) was calculated using the trapezoidal rule, which numerically integrates the luminescence signal over the full duration of the experiment. This approach provides an integrated measure of proliferative output that accounts for early growth kinetics, expansion rate, and saturation behavior, while reducing sensitivity to single time-point fluctuations.

For visualization and comparative analysis of proliferation kinetics, smoothed growth trajectories were generated using generalized additive models (GAM). GAM fitting was applied independently to each cell line and seeding density, modeling luminescence as a smooth function of time. Model predictions were used to derive fitted curves and corresponding 95% confidence intervals, allowing visualization of both central growth trends and variability across replicates. These smoothed profiles were used exclusively for data visualization, while all quantitative comparisons were based on the original normalized measurements and AUC values.

Correlation analyses between proliferative output (AUC) and migration efficiency, expressed by the δ scaling parameter, were performed separately for each seeding density using Pearson’s correlation coefficient.

### Survival Analysis and Stratification Strategy

Retrospective survival analysis was performed on a cohort of 24 patients with IDH-wildtype GBM. Overall survival (OS) was defined as the time from surgical resection to death or last follow-up. Initially, patients were stratified into three groups based on pre-established migration efficiency levels: Low (δ **≤**0.55), Medium (0.55 < δ **≤**0.70), and High (δ>0.70). Kaplan-Meier survival curves were generated, and statistical significance between these cohorts was assessed using the Log-rank (Mantel-Cox) test in GraphPad Prism (Boston, Massachusetts USA). To maximize the prognostic resolution of the δ scaling parameter, a data-driven optimization approach was subsequently employed using Jamovi software (v2.6). An optimal clinical threshold was identified by testing all potential cut-points to find the value that maximized the statistical divergence (log-rank Z score) between groups. This analysis identified a δ scaling cut-off of 0.670, leading to the re-stratification of the cohort into LowC (≤0.670) and HighC (δ > 0.670) populations. The association between migratory efficiency (quantified by the δ scaling parameter) and overall survival was modeled using Cox proportional hazards regression. To account for potential non-linearities in the relationship between cell motility and mortality risk, penalized cubic splines were incorporated into the model. This approach allowed for the estimation of functional forms without assuming a strictly linear effect. Model performance and significance were assessed using the likelihood ratio test and Wald test, while the concordance index (C-index) was calculated to evaluate the model’s predictive accuracy. Hazard ratios (HR) and their corresponding 95% confidence intervals (CI) were derived from the spline fits to visualize the risk transition across the δ spectrum. All spline-based analyses were performed using the survival and rms packages in the R statistical environment.

### Whole Transcriptome Analysis

Raw sequences were generated directly from the NextSeq2000 sequencer using the Dragen BCL convert plugin. The quality of the raw reads was assessed using FastQC (v0.12.1) and potential sample contamination was checked with FastQ Screen (v0.16.0). Reads were subsequently aligned to the hg38 human reference genome, obtained from the UCSC Genome Browser, using STAR aligner (v2.7.9). A raw gene expression count matrix was then generated using feature counts from the Subread package (v2.0.5). All subsequent transcriptomics analyses were performed in R (version 4.4.0). To identify potential genes underlying the different cell movement phenotypes, we performed differential expression (DE) analysis between the patient groups identified in the phenotypic analysis. The DE analysis was conducted using the DESeq2 package^41^. To reduce the effect of genes with low counts, the estimated values were shrunken using the ‘normal’ shrinkage method.

### Gene Set Enrichment Analysis (GSEA)

Following this correction, genes were considered differentially expressed if they met the criteria of an adjusted p-value < 0.05 and an absolute value of the log2fold-change > 0.58 (corresponding to a fold-change of 1.5). A GSEA was also performed on the results of the differential expression analysis. GSEA was executed querying the Gene Ontology Biological Processes database through the gseGO function of clusterProfiler R library^18^.

### Whole exome analysis

Genomic DNA was extracted from the original fresh tissue samples using Maxwell 16 Tissue LEV DNA Purification Kit (Promega, Madison, WI, USA), following the manufacturer’s instructions. Whole-exome library preparation was performed using Illumina DNA Prep with Exome 2.5 Enrichment (Illumina, San Diego, CA, USA), following manufacturer’s procedure. Paired-end sequencing was performed using NextSeq 2000 (Illumina, San Diego, CA, USA) with 101 bp of read length loading on NextSeq 1000/2000 P2 Reagents (200 Cycles) v3. WES was also performed on the patients’ blood to subtract germline mutations.

After germline subtraction, somatic variants were annotated using CRAVAT^42^ and filtered based on population frequency (<0.01%), coding impact (frameshift, missense, stop-gain), and sequencing quality metrics (DP ≥ 30, MBQ/MMQ/GERMQ ≥ 20, ECNT ≤ 2, ROQ ≥ 20, NALOD > 0, FS ≤ 20, SOR ≤ 3) to exclude potential artifacts.

### Tree decision model approach

#### Data pre-processing and filtering

Raw data were subjected to batch effect correction, normalization, and feature selection. Given the lack of a consensus on the optimal preprocessing pipeline, several alternatives were evaluated using an exhaustive grid-search approach.

For batch effect correction, ComBat-seq^43^ was employed, specifying the “delta class (high/mid/low)” as a covariate. Normalization was performed using the Python library included in the DeClut software^44^. Several methods were tested: Counts Per Million (CPM), Reads Per Kilobase Million (RPKM), Transcripts Per Million (TPM), Trimmed Mean of M-values (TMM), Median Quartile (MQ), Upper Quartile (UQ), and Mean of Ratios (MOR).

Feature selection was conducted by partitioning genes into 50 clusters using k-means. The average count value for each centroid was calculated, and clusters were iteratively removed in increasing order of expression until a target number of genes was reached. Multiple target thresholds were tested (500, 1,000, 2,000, 5,000, 7,000, and 10,000 genes), as well as a configuration where only the cluster with the minimum average count was removed.

To evaluate preprocessing quality, the Mean Squared Error (MSE) of a Support Vector Regressor (SVR), trained to predict the delta values of the samples, was used. SVR hyperparameters were optimized via exhaustive grid-search, testing four kernels (linear, poly, RBF, sigmoid) and six values for the regularization parameter C = (0.001, 0.01, 0.1, 1, 10, 100). Due to the small sample size, a leave-one-out (LOO) cross-validation scheme was adopted. The pipeline was implemented in Python (v.3.13.7) using the scikit-learn (v.1.8). The results indicated that batch-effect correction was overly aggressive, generally increasing the MSE; consequently, it was omitted from the final pipeline. The MOR normalization strategy yielded the most consistent distribution of library sizes, while the removal of a single cluster proved most effective in minimizing the MSE.

#### Raw data have AI-based identification of genes involved in delta

An SVR model was utilized to compile a list of genes correlated with the delta variable. The model was trained on the entire dataset. Since the objective was model explainability rather than the creation of a generalizable predictor, the potential bias or overfitting associated with using the same samples for training and testing was considered acceptable for feature importance extraction. For each sample, the SHAP library^45^ was employed to identify relevant genes (those with a SHAP score > 0). A per-sample contribution score, proportional to the Jaccard similarity between the delta values and the model’s predictions, was further calculated. The final per-gene score was computed as the sum of these contributions across all samples where the gene was deemed relevant. This procedure yielded a refined list of 34 candidate genes. To develop an explainable AI model, this gene list was used to train a Decision Tree. Given the limited sample size, a leave-one-out strategy was again employed. To account for structural instability—where the inclusion/exclusion of a single sample significantly altered the tree—the gene list was further narrowed to only those features selected at least once during the LOO iterations. Finally, a Random Forest ensemble, consisting of three decision trees, was trained to stabilize the findings

