## Supplementary material for "From Single-Cell Emergent Behaviors to Clinical Outcome: PTEN-driven Migratory Efficiency as a Potential New Vulnerability in Glioblastoma": Suppl Figures

### Supplementary Material Morelli et al 2026

### Supplementary VideoS1-S2

Video S1.avi

Video S2.mp4

**Video S1. Representative video:**Phase-Contrast (Raw): Time-lapse acquisition (10x magnification, 1 frame/5 min) showing primary GBM cell motility over 18 hours in glass-bottom dishes.

**Video S2. Representative video with Segmentation & Trajectories:** Computational analysis of the raw footage. Cell nuclei/bodies were segmented via TrackMate-Cellpose integration and tracked using the LAP algorithm. Each distinct color identifies an individual cell trajectory, utilized to quantify emergent migratory behaviors and dynamic traits.

### Figure S1.

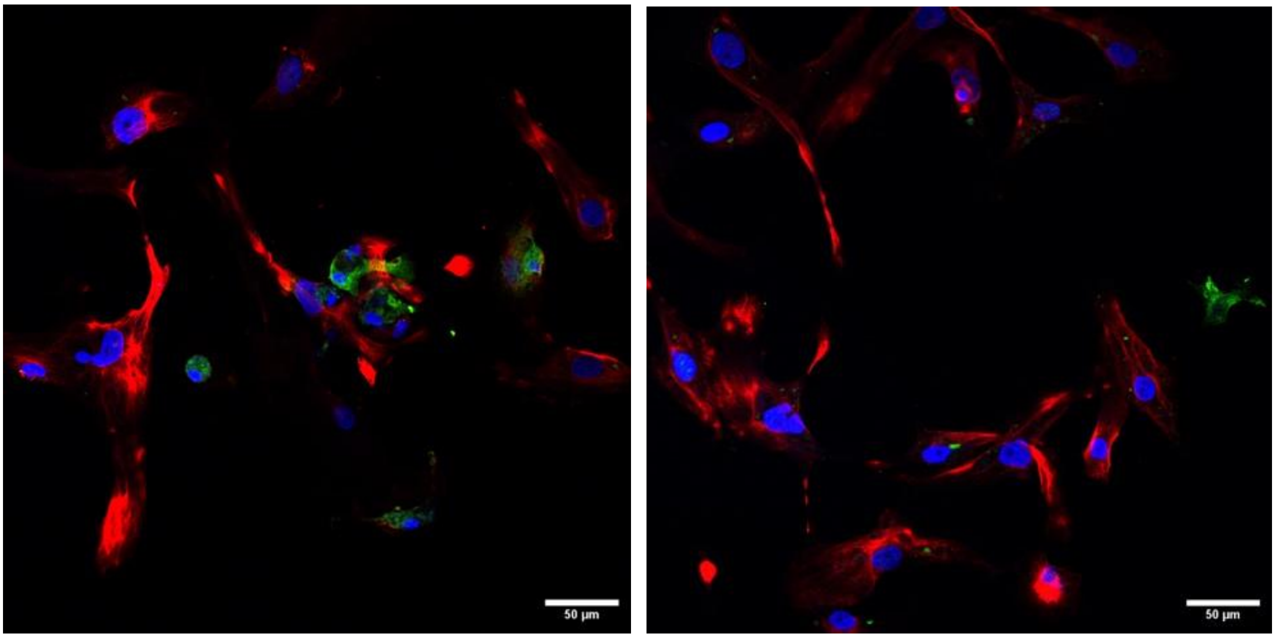


**Figure S1. Immunofluorescence characterization of patient-derived primary glioblastoma (GBM) cultures.** Two representative confocal photos showing the cellular composition of a primary GBM culture. Cells were immunostained for GFAP (red), marker for malignant glial cells and astrocytes; IBA1 (green), identifying tumor-associated microglia/macrophages (TAMs); and DAPI (blue), providing nuclear counterstaining. The image highlights the spatial distribution and morphological diversity of the tumor and immune cell populations. Scale bar: 50 µm.

### Figure S2

#### Colony Formation and Growth Dynamics

To contextualize our single-cell migration analysis, we evaluated the long-term proliferative behavior of patient-derived primary GBM cell cultures via colony formation assays. While commercial glioblastoma lines readily formed dense, quantifiable clusters, primary GBM cell cultures demonstrated a markedly reduced capacity for colony formation across all δ scaling groups. Instead, these cells remained largely dispersed, exhibiting a preference for individual growth and migration over collective aggregation. This observation provides a critical biological rationale for our study's methodology: the lack of traditional colony-forming behavior confirms that glioblastoma primary cells are best characterized through single-cell behavioral analysis rather than bulk population metrics. Importantly, we note that while immortalized commercial cell lines consistently form dense colonies, a behavior mirrored even by Normal Human Astrocytes (NHA), patient-derived primary cells deviate from this pattern, favoring a more infiltrative, individualistic growth phenotype (**Figure S2**). This distinction suggests that established commercial lines may have lost the characteristic single-cell dynamics inherent to glioblastoma during the immortalization process. Consequently, our findings demonstrate that patient-derived primary models provide a significantly more reliable and physiologically relevant platform for studying the discrete migratory patterns and emergent complexity that define GBM progression.

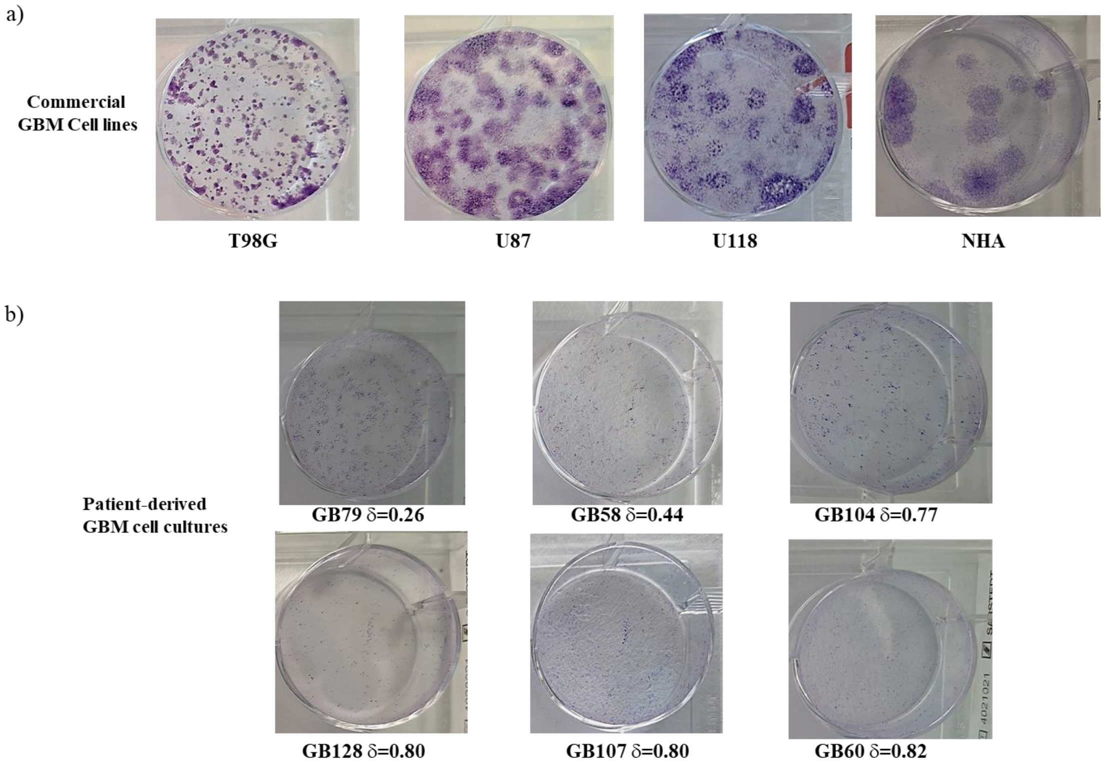


**Figure S2. Comparative Colony Formation Assay between Commercial Cell Lines and Patient-Derived GBM Primary Cultures. (a)** Commercial Cell Lines: Representative images of colony formation assays for established glioblastoma lines (T98G, U87, U118) and Normal Human Astrocytes (NHA). These lines demonstrate a high capacity for anchorage-independent growth, forming dense and quantifiable macroscopic clusters. **(b)** Patient-Derived GBM Cell Cultures: Representative images of primary cultures organized by their respective δ values: GB1 (δ = 0.27), GB4 (δ = 0.45), GB19 (δ = 0.66), GB25 (δ = 0.74), GB26 (δ Delta = 0.75), and GB27 (δ Delta = 0.78). In contrast to commercial models and normal control astrocytes, primary cells exhibit a significantly reduced ability to form colonies regardless of their δ group, predominantly existing as dispersed, single cells.
